# Integrative genomics of *Plasmodium knowlesi* reveals parasite-intrinsic regulators of severe human malaria

**DOI:** 10.64898/2026.03.22.713451

**Authors:** Jacob A F Westaway, Ernest Diez Benavente, Michal Kucharski, Sourav Nayak, Duong T Q Huy, Sarah Auburn, Timothy William, Giri S Rajahram, Kim A Piera, Hidayat Trimarsanto, Kian Soon Hoon, Caitlin Bourke, Anisah Jantim, Abdul Marsudi Manah, William Gotulis, Robert W Moon, Roberto Amato, Bridget E Barber, Chris Drakeley, Nicholas M Anstey, Matt A Field, Zbynek Bozdech, Matthew J Grigg

## Abstract

To dissect parasite determinants of clinical disease severity in *Plasmodium knowlesi*, we performed the first integrative genome-wide association (GWAS), transcriptomic, and expression quantitative trait locus (eQTL) analysis from clinical malaria isolates. We identify distinct parasite programs associated with WHO-defined severe disease, characterized by transcriptional activation of stress-response and host–interaction pathways. In contrast, invasion-linked programs were more strongly associated with parasite burden, while immune-evasion pathways showed overlapping but distinct associations with both burden and severity. GWAS and eQTL analyses revealed genetic regulation of transcriptional states, including immune-evasion variant antigen families (*SICAvar* and *kir*) and chromatin-associated regulators. Mature gametocyte transcripts were detectable across infections, including those with low parasitaemia, indicating that transmission-stage expression occurs independently of both clinical severity and parasite density. Together, these findings show that severe *P. knowlesi* malaria is associated with genetically regulated parasite transcriptional programs that are not fully explained by parasite burden.

## Main

*Plasmodium knowlesi*, a zoonotic malaria parasite naturally infecting long-tailed and pig-tailed macaques, has emerged over the past two decades as the dominant cause of malaria in Malaysia and represents an increasing threat across Southeast Asia (1, 2, 3). Unlike *P. falciparum* and *P. vivax*, whose incidence has declined under sustained control efforts, *P. knowlesi* is increasing due to land-use change that intensifies human exposure to macaque reservoirs and *Anopheles* vectors (4, 5, 6). The zoonotic transmission of *P. knowlesi*, involving intractable macaque reservoirs and outdoor forest-edge exposure, poses a fundamental barrier to elimination (2, 7, 8). As a result, *P. knowlesi* is now recognized as a major obstacle to eliminating malaria in the broader region (9, 10).

Human *P. knowlesi* infections exhibit a broad clinical spectrum, ranging from asymptomatic parasitaemia to uncomplicated febrile illness and severe, life-threatening disease. In endemic settings, adult case fatality in adults are comparable to those seen with *P. falciparum* (11, 12). Severe manifestations include acute kidney injury, jaundice, and respiratory distress, and their risk increases with both age and parasite density (11, 12). Strikingly, fatal outcomes can occur even at modest parasitaemia (13), suggesting that in addition to excessive host immune responses, *P. knowlesi*-specific factors beyond biomass and outside of red blood cell invasion mechanisms may contribute to pathology (14). Contributing host determinants of both infection risk and severe disease have been described, including a well-documented higher risk of infection among adult males, largely attributed to behavioural and occupational exposure patterns (7, 15). However, the role of parasite genetics and transcriptional programs in modulating parasitaemia and clinical severity remains poorly understood. Identifying these parasite-intrinsic drivers is therefore critical to explain disease heterogeneity and elucidate how a zoonotic parasite may have adapted to cause severe clinical outcomes in humans.

Genomic studies of *P. knowlesi* to date have primarily focused on population structure, zoonotic transmission, and signatures of selection, providing insight into its emergence as a major human pathogen (16, 17, 18), but offering limited understanding of mechanisms underlying clinical severity. These analyses have identified host-associated subpopulations and selective sweeps in genes linked to invasion and antigenic variation, suggesting that adaptation to macaque—and potentially human—hosts has shaped parasite diversity. Functional genomic studies have identified factors critical for host interaction; for example, *Moon et al.* (2016) demonstrated that the invasion ligand NBPXa, a reticulocyte-binding protein family member, is essential for erythrocyte invasion in humans (19). Whether parasite-intrinsic genetic variation contributes directly to clinical severity—independent of host immunity or parasite biomass—remains unclear. In *P. falciparum*, specific var gene expression profiles are strongly associated with severe disease (20), demonstrating that parasite genetic and regulatory variation can influence clinical outcome.

In contrast, transcriptomic investigations of *P. knowlesi* remain limited. While the intraerythrocytic developmental cycle (IDC) has been extensively mapped in *P. falciparum* and *P. vivax*, stage-specific expression and transcriptional programs in *P. knowlesi* are poorly characterized (21, 22), and their relevance to human disease unclear. Moreover, the genetics and regulation of multigene families such as schizont-infected cell agglutination variant (*SICAvar*) genes and knowlesi-interspersed repeat (*kir*) remain difficult to characterise, as their sequence diversity hampers both genomic and transcriptional analyses (23, 24, 25, 26). Although regulatory variation in such multigene families is increasingly recognized as a major driver of parasite biology and pathogenesis in other *Plasmodium* species (27, 28, 29, 30, 31, 32, 33), they have not been systematically studied in *P. knowlesi*. To date, no integrated genome–transcriptome analysis has examined how parasite genetic and transcriptional variation interact to influence clinical severity in *Plasmodium* species.

To address these knowledge and methodological gaps, we performed an extensive integrative genomic and transcriptomic analysis of clinical isolates of *P. knowlesi* malaria. Using genome-wide association studies (GWAS), we identified parasite genetic variants associated with clinical severity and parasitaemia. In parallel, we generated a *de novo* transcriptome assembly to capture both conserved and highly divergent gene families, applying a hybrid annotation strategy that combined reference-based mapping with homology- and domain-informed annotation. This enabled robust identification of differentially expressed genes (DEGs) and pathway-level shifts through Gene Ontology (GO) enrichment. We further leveraged IDC deconvolution to disentangle stage composition effects from transcriptional signatures, enabling assessment of parasite synchrony potential contribution of stage composition to disease severity. Finally, we performed expression quantitative trait locus (eQTL) mapping to link parasite genotypes to transcriptional regulation *in vivo*. Together, this work represents the first parasite genome–transcriptome association study in *P. knowlesi* parasites infecting humans, providing a comprehensive framework able to dissect how parasite variation influences replication dynamics and clinical outcome.

## Results

### Distinct genetic architectures of disease severity and parasite burden converge on growth and invasion pathways

We performed genome-wide association studies (GWAS) to investigate the genetic basis of *P. knowlesi* infection outcomes. Whole-genome sequencing of parasite DNA from 737 patients in Sabah, Malaysia, followed by stringent variant and sample filtering, including restriction to infections classified as clonal (F_ws_ ≥95; 88% of samples), yielded 77,137 high-quality SNPs across 424 samples. ADMIXTURE, neighbour-joining and principal components analyses (PCA) confirmed three established major genetic clusters (*Mf* and *Mn*, associated with *Macaca fascicularis* and *Macaca nemestrina*, respectively, and a *Peninsular* Malaysia lineage) (16) (Figure 1A-B).

**Figure 1.**
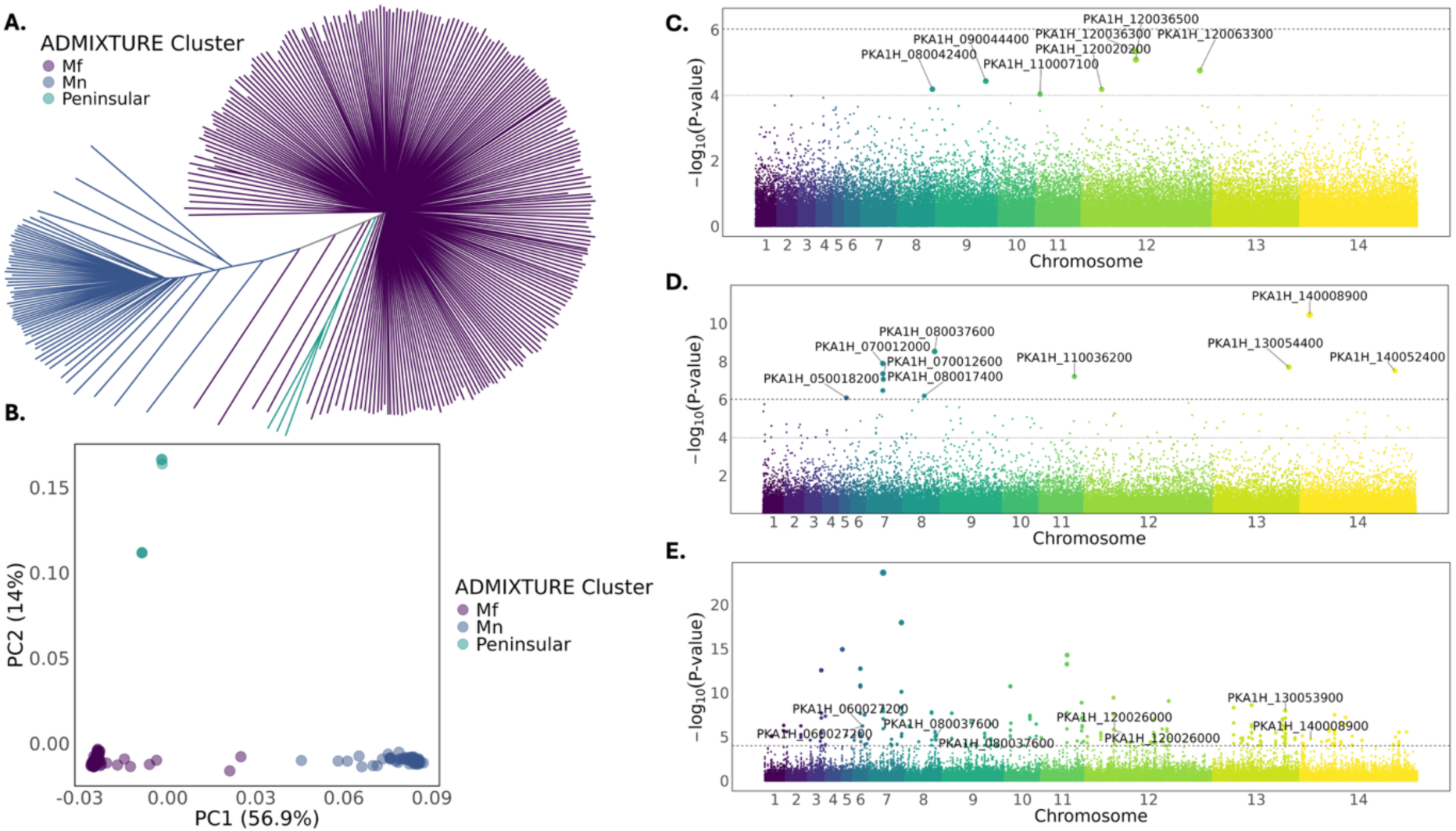
Genome-wide association studies of *Plasmodium knowlesi* and population structure. **(A)** Population structure of *P. knowlesi* clinical isolates inferred by ADMIXTURE clustering, visualized as a neighbour-joining tree, with isolates coloured by major subpopulation (*Mf*, *Mn*, *Peninsular*). **(B)** Principal component analysis (PCA) plot of *P. knowlesi* isolates, coloured by ADMIXTURE cluster (Supplementary Figure 1). **(C)** Manhattan plot of the disease severity-based GWAS. The upper dashed line indicates the genome-wide significance threshold after Bonferroni correction and the lower dotted line indicates a suggestive threshold adjusted for genome size (p < 1×10⁻⁴), with suggestive hits labelled with PKA1H1 gene identification. **(D)** Manhattan plot of the disease parasitaemia-based GWAS. The upper dashed line indicates the genome-wide significance threshold after Bonferroni correction and the lower dotted line indicates a suggestive threshold adjusted for genome size (p < 1×10⁻⁴), with significant hits labelled with PKA1H1 gene identification. **(E)** Manhattan plot of cross-population extended haplotype homozygosity (XP-EHH) analysis with significant selection threshold of (p < 1×10⁻⁴) represented by the dotted line and gene hits overlapping GWAS labelled with PKA1H1 gene identification.

Next, we examined clinical disease severity given its public health relevance, followed by parasitaemia as a quantitative trait reflecting parasite-intrinsic growth dynamics. Associations were tested for WHO-defined severe malaria (34) (severe n = 111, uncomplicated n = 313; Supplementary Table 1) and microscopy-quantified parasitaemia (median 9.07 × 10³ parasites/µL, IQR 4.36 × 10³–2.64 × 10⁴). To identify putative genetic effector of both clinical phenotypes, we carried out GWAS with the generated WGS dataset, with a linear mixed model incorporating a genetic relationship matrix (GRM) to control for relatedness and fine-scale population structure. Across both traits, 121 SNPs surpassed a suggestive threshold of p < 1×10⁻⁴ (Figure 1C–E), including 11 loci reaching genome-wide significance for parasitaemia after Bonferroni correction (Figure 1C; Figure 1D; Supplementary File 2). Minimal overlap was observed between severity- and parasitaemia-associated loci, indicating largely distinct parasite genetic architectures for the two phenotypes. Cross referencing the annotated GWAS-implicated genes with the transposon insertion datasets generated by *Oberstaller et a.l* (35, 36), revealed that several loci associated with both parasitaemia and disease severity include both genes essential for asexual blood-stage growth and genes dispensable under laboratory conditions (Supplementary File 2).

Using the suggestive statistical threshold (p < 1×10⁻⁴), we identified seven distinct parasite loci located on chromosomes 8, 9, 11 and 12 associated with disease severity (Figure 1C). There were three SNPs on chromosome 12 linking S-adenosylmethionine–dependent methyltransferase (PKA1H_120036300), a zinc finger protein (PKA1H_120036500) and phosphoenolpyruvate carboxykinase (PKA1H_120063300), and a potassium channel (PKA1H_120020200) with severity of *P. knowlesi* infection within this cohort. Notably, phosphoenolpyruvate carboxykinase links central carbon metabolism to parasite growth (37), while the S-adenosylmethionine–dependent methyltransferase implicates chromatin-mediated regulation (38). SNPs on chromosomes 8, 9 and 11 mapped to genes annotated as *Plasmodium proteins, unknown function*. In addition to the GWAS-based loci, we explored functional assignments of genes within the top-ranked severity-associated SNPs (top 100 by p-value). Here we noted several invasion-and export-associated genes, including an MSP7-like protein (PKA1H_120070500), MSP5 (PKA1H_040019100), and a PHIST protein (PKA1H_060028600), alongside transcriptional regulators such as an α/β-hydrolase domain protein (PKA1H_020018900) and a CCR4–NOT complex subunit (PKA1H_090006900) (Supplementary File 2). Taken together, these results suggest roles for transcriptional regulation, metabolic adaptation, and ion transport, as well as processes related to invasion and host-cell remodelling, which contribute to severe outcomes of *P. knowlesi* infections in human zoonotic infections.

To identify genetic factors underlying parasite load, we carried out a GWAS with peripheral parasitaemia as a quantitative trait. Here we identified eleven genome-wide significant SNPs (Bonferroni corrected p < 9.63×10^-7^) including a putative peptidase (PKA1H_140008900), a DEAD-box helicase (PKA1H_080037600), a cyclin-related protein (PKA1H_140052400), and a zinc finger protein (PKA1H_050018200), with orthologues implicated in cell-cycle progression, chromatin-associated transcriptional regulation, and RNA metabolism and development in *P. falciparum* (39, 40, 41, 42). A prominent association peak on chromosome 7 mapped to several conserved proteins of unknown function, including PKA1H_070012000 and PKA1H_070012600. Structural modelling using AlphaFold2 revealed multiple high-confidence folded domains for PKA1H_070012000, and FoldSeek analysis identified structural homology to a lipid-binding protein (GO:0008289; Supplementary Figures 3-4; Supplementary Table 2), suggesting potential roles in membrane-associated or export processes. Similar to the severity analysis (above), additional suggestive signals (p < 1×10⁻⁴) included five SNPs on chromosome 7 mapping to a dynein heavy chain gene (PKA1H_070007900) (Supplementary File 2). Independent signals were also observed in two additional dynein heavy chain genes located on chromosomes 2 (PKA1H_020020400) and 13 (PKA1H_130007100), as well as in a dynein light intermediate chain gene on chromosome 7 (PKA1H_070009100) and several loci close to myosin light chain B (PKA1H_090022200) and kinesin-4 (PKA1H_080014400). These observations collectively link the dynein motor complex and other motility-related factors with *P. knowlesi* parasite burden. We also identified SNP variants linked with DNA2/nam7 helicase (PKA1H_090009800), zinc finger protein (PKA1H_050018200), CCR4-associated factor 1 (PKA1H_140034600), and tyrosine kinase-like protein (PKA1H_120056200). These additional loci complement the cyclin- and DEAD-box–associated signals, reinforcing roles for cell-cycle–linked transcriptional control and RNA metabolism, while independently implicating cytoskeletal motor function and kinase-mediated signalling pathways involved in invasion, egress, and stage-specific development (42, 43, 44, 45). This functional footprint appears highly distinct from the severity-linked genetic factors that are implicated in host-parasite interactions and metabolic adaptations.

To assess whether GWAS-implicated loci show signatures of recent adaptive evolution, we performed cross-population extended haplotype homozygosity (XP-EHH) analyses comparing parasites from severe and uncomplicated infections (Figure 1E; Supplementary File 2). Several strong XP-EHH selection haplotypes were observed at multiple members of the dynein motor complex, including two dynein heavy chains (PKA1H_030018000; PKA1H_060012500), with haplotypes enriched among parasites from uncomplicated infections. Although these loci are distinct from the dynein-implicated variants identified by GWAS, they suggest pathway-level convergence on cytoskeletal function under recent selective pressures. XP-EHH signals were also detected at two loci associated with greater parasitaemia in the GWAS, including the putative peptidase (PKA1H_140008900), enriched in severe infections, and the DEAD-box helicase (PKA1H_080037600), enriched in uncomplicated infections. Given the zoonotic transmission ecology of *P. knowlesi*, these selection signals likely reflect ongoing adaptive dynamics within circulating parasite populations rather than lineage-specific fitness advantages uniquely supporting severe human infection in Sabah.

### Differential gene expression across the clinical spectrum of disease severity reveals divergent invasion- and regulatory-associated programs, independent of intraerythrocytic life-stage composition

We carried out RNA sequencing on 210 clinical blood samples that passed stringent RNA quality control thresholds (including RIN ≥4 and purity metrics; see Methods), following initial extraction from 402 field specimens, including 39 from patients with severe malaria and 171 from uncomplicated cases. Following rRNA depletion and Illumina library preparation, as previously described for *P. falciparum* (46), reads were quality filtered and human-derived sequences removed. To capture both conserved and highly variable parasite transcripts, we constructed a *de novo* meta-transcriptome using Trinity (47), allowing recovery of hypervariable multigene families such as *SICAvar* and *kir*. The assembly contained ∼4.7 million transcripts, with an Ex90 of 65,634 and an Ex90N50 of 5.5 kb, indicating high contiguity among the most abundantly expressed transcripts. After stringent filtering for expression and quality, 42,033 transcripts were retained for downstream analyses (Supplementary Tables 3-4). Transcripts were functionally annotated using a hybrid strategy (Supplementary Figure 6) that combined reference-based mapping to the *P. knowlesi* PKA1H1 genome (48) with *de novo* annotation, enabling comprehensive assignment of coding, non-coding, and structurally complex features (49). Expression counts were calculated by mapping reads back to the transcriptome assembly.

As a first step, we performed transcriptome deconvolution using Scaden (50) to estimate parasite life-stage composition using the reference panels derived from single-cell data for *P. knowlesi* and *P. berghei* from the Malaria Cell Atlas Project (21), with *P. berghei* used for its similar 24-hour asexual cycle and early gametocyte maturation kinetics (21). Deconvolution revealed heterogeneity in IDC composition across infections (Figure 2A), with trophozoites predominating (51). Gametocyte transcripts comprised approximately 25% of the inferred IDC and did not differ between severe and uncomplicated infections, nor were they associated with peripheral parasitemia (Supplementary Table 5; Supplementary Figure 7). Gametocytes were detectable even at very low parasitaemia, including below routine microscopy detection thresholds (Figure 2C; Supplementary Table 7). Elevated peripheral schizont proportions have been proposed as a marker of severe *P. knowlesi* malaria (52); however, in this cohort we found median schizont transcript proportions (2.7% [IQR 1.8–6.3%]) were similar to those with uncomplicated infections (3.5% [IQR 1.9–8.2%]). Likewise, the proportion of infections with

**Figure 2.**
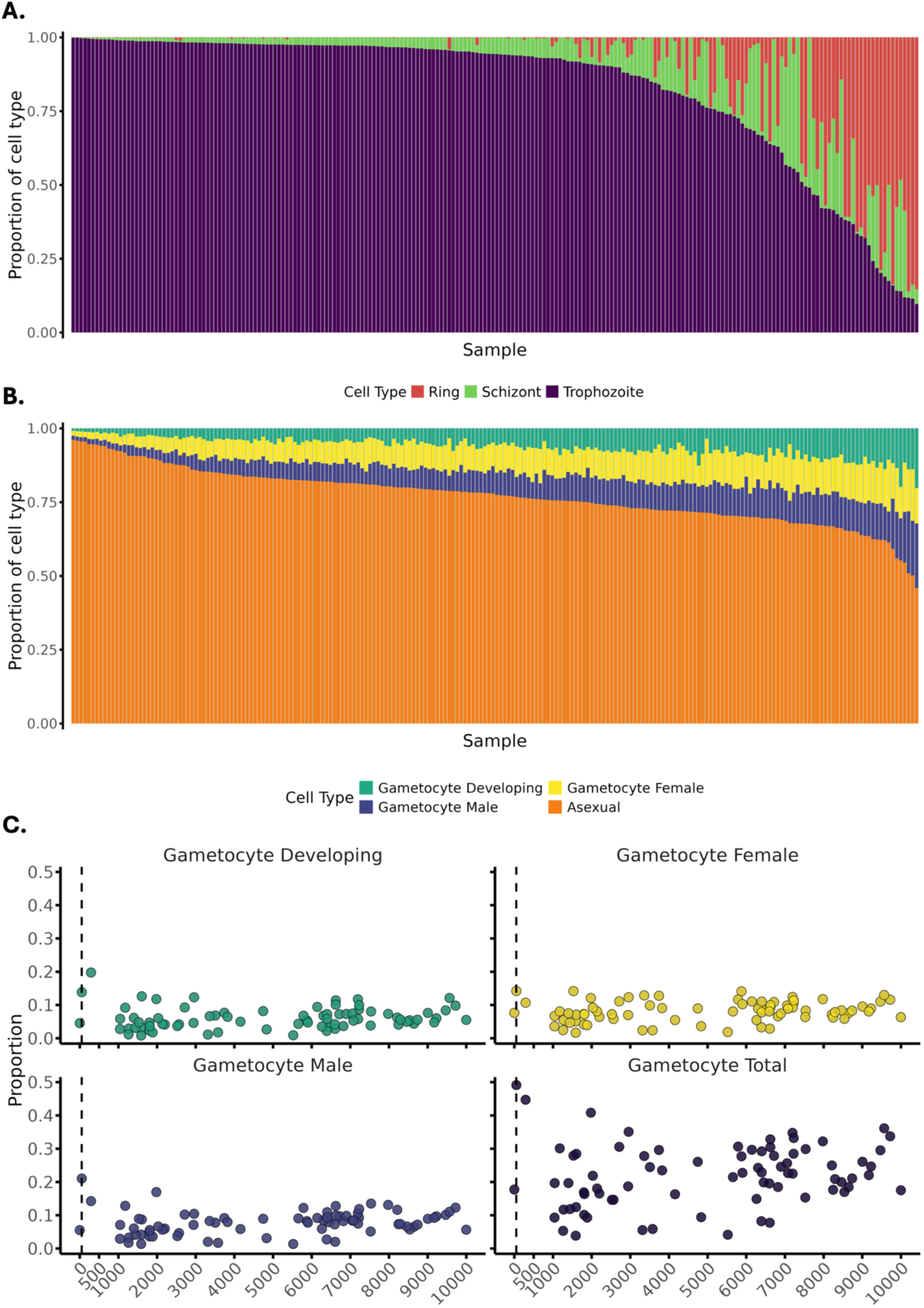
Intraerythrocytic developmental cycle (IDC) composition of *Plasmodium knowlesi* infections. Proportions of asexual life stages across individual infections estimated with Scaden using *P. knowlesi* **(A)** and *P. berghei* **(B)** single cell reference sets. **(C)** Scatter plots of Scaden-predicted proportions for developing, female, male, and total gametocytes as a function of parasitemia (subset to <10000 parasites/µL) for individual clinical isolates. Dashed vertical line represents the limit of routine microscopy detection (50 parasites/ µL). Predictions were generated using a *Plasmodium berghei* single-cell reference. Points represent individual samples and are coloured by level of parasitemia.

>10% schizont transcripts, including after adjustment for parasitaemia, did not differ between groups. Together, these findings indicate that large-scale differences in IDC composition does not account for severity-associated transcriptional differences.

To investigate parasite transcriptional correlates of clinical outcome, we performed differential expression analysis using a consensus framework (Supplementary Figure 8), comparing isolates from patients with severe versus uncomplicated malaria. Exploratory PCA revealed no discrete clustering by severity (Figure 3A), although PC1 scores difference significantly between clinical groups (Figure 3B, Wilcoxon *p* < 0.0001), indicating that disease severity contributes to global expression variation. Assessment of potential confounders showed that technical variables, including library size, mapped read counts, transcripts detected, and IDC composition generally explained little variance (R² < 0.14), with the exception of transcripts detected in PC3 (p < 0.001, R² = 0.49; PC3 = 2.51%) (Supplementary Tables 8–9).

**Figure 3.**
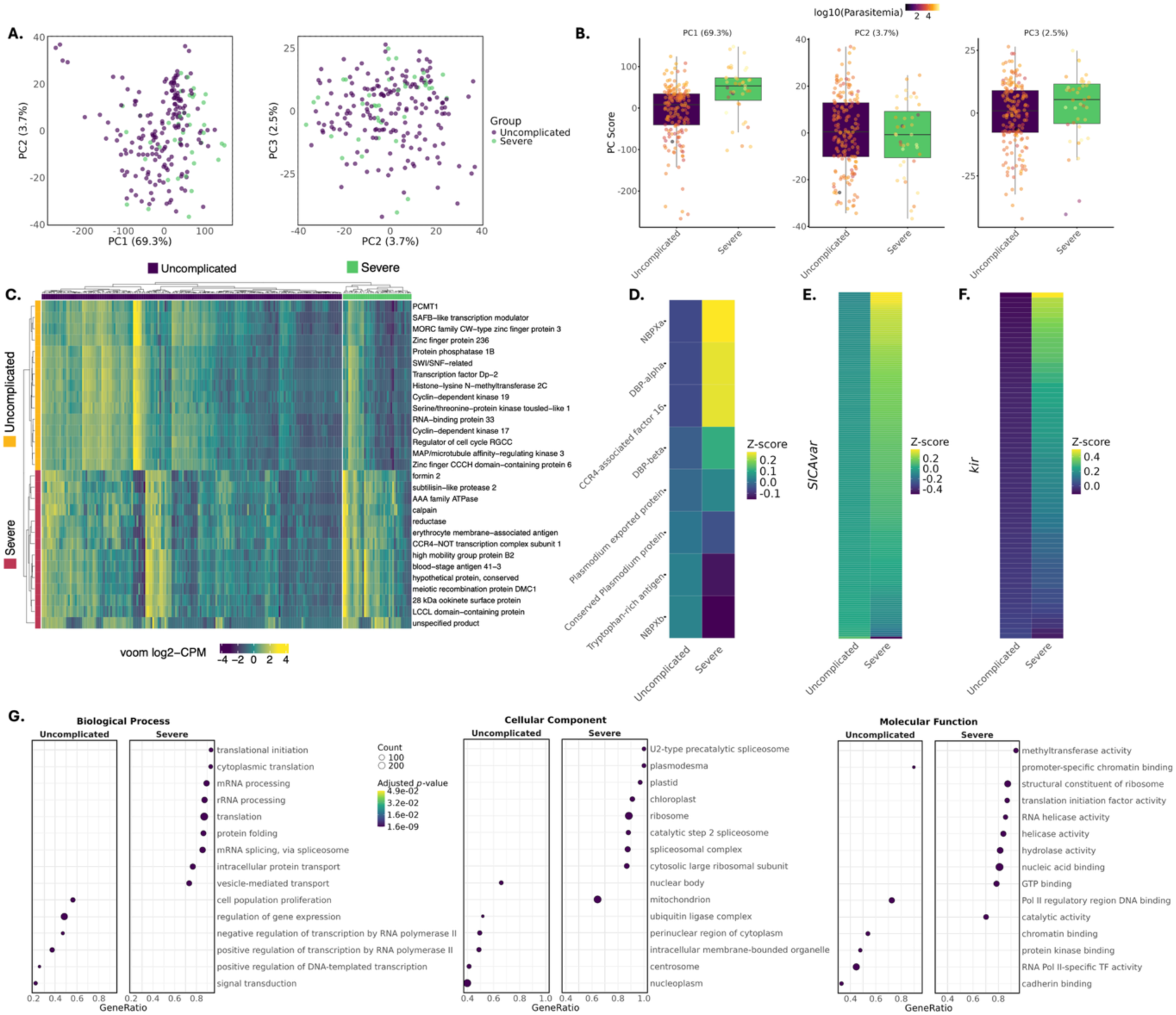
Comparative transcriptomic analyses of *P. knowlesi* infections associated with severe and uncomplicated malaria. **(A)** Principal component analysis (PCA) of voom-transformed, length-filtered, parasite transcriptomes, showing relationships among samples based on the leading components of expression variation. **(B)** Boxplots of sample scores for the top principal components stratified by clinical severity (Sm, severe malaria; Um, uncomplicated malaria), with individual samples coloured by log-transformed parasitaemia. **(C)** Heatmap of row-scaled (z-score) voom-normalised expression values scaled within the displayed gene set for the 15 parasite genes most strongly upregulated in severe or uncomplicated infections. Samples are grouped by clinical severity and hierarchically clustered within groups; rows are clustered by expression similarity. Side annotations indicate the clinical group in which each protein shows higher expression. Full list of differentially expressed transcripts can be found in Supplementary File 3. Abbreviations: PCMT1, protein-L-isoaspartate O-methyltransferase 1; SWI/SNF-related, SWI/SNF-related matrix-associated actin-dependent regulator of chromatin subfamily A member 5. **(D)** Group-level z-score summaries of host-adaptation associated genes identified from long-term growth in human vs macaque erythrocytes (Moon et al., 2016); **(E)** the *SICAvar* multigene family; **(F)** the *kir* gene family: highlighting contrasting expression patterns between severe and uncomplicated infections. **(G)** Dot plots showing the top 15 enriched Gene Ontology (GO) terms within each ontology (Biological Process, Cellular Component, and Molecular Function). Terms are ordered by GeneRatio, with point size indicating the number of genes contributing to each term and colour representing the adjusted *p*-value. For readability, two GO terms were abbreviated in panel G: *Pol II regulatory region DNA binding* (RNA polymerase II transcription regulatory region sequence-specific DNA binding) and *RNA Pol II-specific TF activity* (DNA-binding transcription factor activity, RNA polymerase II-specific).

Differential expression analysis identified 129 transcripts whose mRNA abundance differed significantly between severe and uncomplicated infections (FDR < 0.05). Heatmap inspection revealed greater within-group consistency among transcripts upregulated in severe malaria, whereas uncomplicated infections showed increased inter-sample variability, suggesting more heterogeneous transcriptional states (Figure 3C). Differentially expressed transcripts included conserved parasite genes, regulatory factors, and variant antigen families (Figure 3C; Supplementary File 3; Supplementary Table 10), with 30 transcripts also associated with parasitaemia in an independent model (Supplementary Table 11). Several severity-upregulated transcripts also overlapped GWAS signals, supporting their potential relevance to disease biology. Among the top differentially expressed genes was CCR4–NOT transcription complex subunit 1, consistent with CCR4-related biology identified by GWAS (Supplementary File 2). CCR4–NOT complexes function as conserved post-transcriptional regulators in *Plasmodium*, coordinating invasion and egress protein expression as well as broader stage-specific transcriptional programs (44, 53). Additional upregulated genes included subtilisin-like protease 2, whose orthologue in *P. falciparum* is required for merozoite egress and invasion (54), and high mobility group protein B2 (HMGB2), a chromatin-associated factor implicated in inflammatory responses during experimental cerebral malaria in *P. berghei* (55). Transcripts encoding host–parasite interface proteins were also enriched, including blood-stage antigen 41-3 and erythrocyte membrane-associated antigens, consistent with altered host cell interaction during severe infection. A smaller subset of genes associated with gametocyte or transmission-stage development, including DMC1 (56), LCCL domain-containing protein (57), and a 28 kDa ookinete surface protein (58), was also upregulated. Collectively, these results suggest that *P. knowlesi* in severe malaria mount a transcriptional program geared towards virulence, host–parasite interaction, and stress adaptation.

Analysis incorporating parasitaemia as a continuous covariate identified several transcripts that overlap with severity-associated transcriptional signals, including a 28 kDa ookinete surface protein, a meiotic recombination protein, blood-stage antigen 41-3 and erythrocyte membrane–associated antigens (Supplementary File 4). *SICAvar* transcripts comprised a substantial fraction of parasitaemia-associated genes, consistent with broader activation of variant antigen repertoires at higher parasite biomass. Parasitaemia was additionally associated with increased expression of chromatin-related genes, including histone transcripts such as histone H3. Although chromatin-associated genes did not directly overlap with severity-associated transcripts, their enrichment suggests convergence between parasite biomass and severity-linked transcriptional programs, consistent with the collinearity between these traits (Supplementary Table 13). Most severity-associated signals remained independent of parasitaemia, indicating that severe disease involves broader transcriptional changes beyond those attributable solely to parasite load.

### Coordinated activation of invasion ligands, variant antigen repertoires, and stress-associated transcriptional programs in severe malaria

In addition to global analyses, we performed targeted expression analyses of parasite gene families and loci previously implicated in host-specific erythrocyte interactions and adaptation by *Moon et al.* (19). Genes preferentially retained in human-adapted parasites, including NBPXa and CCR4-associated factor 16, showed higher expression in severe malaria, whereas genes linked to macaque-adapted or slower-growing phenotypes, such as NBPXb, were enriched in uncomplicated infections (Figure 3D) (59). Notably, CCR4-associated factor 16 is dispensable during prolonged growth in macaque erythrocytes. *SICAvar* and *kir* gene families also showed higher overall expression in severe malaria, despite substantial heterogeneity among individual transcripts (Figure 3C; Supplementary Figure 9). At the transcriptome-wide level, invasion-associated genes, *SICAvar*, and *kir* transcripts occupied higher expression ranks in severe relative to uncomplicated infections (Supplementary Figure 10). Although few individual *SICAvar* or *kir* transcripts reached transcript-level FDR significance, severe malaria cases showed broader variant antigen repertoire activation, including a significantly higher number of expressed *kir* transcripts per infection compared to uncomplicated cases (median difference = 6.5 transcripts; Wilcoxon *p* = 0.002; Supplementary Figure 11). The single *SICAvar* domain transcript upregulated in uncomplicated malaria likely reflects repertoire composition differences rather than contradiction of the broader family-level pattern. Together, these results indicate coordinated activation of invasion and variant antigen gene families in severe malaria, consistent with broader family-level transcriptional shifts rather than isolated transcript-specific effects.

We performed Gene Ontology (GO) enrichment analysis to detect pathway-level shifts not evident at the single-gene level. In severe *P. knowlesi* malaria, GO enrichment highlighted vesicle-mediated transport and vacuolar membrane categories (Supplementary File 5), together with translational machinery, protein folding, and RNA processing pathways (Figure 3G). These enrichments indicate increased biosynthetic and trafficking activity in parasites from severe infections. Categories related to RNA processing and spliceosome function are broad but have been implicated in regulatory processes controlling clonally variant gene families in other *Plasmodium* species; however, our analysis does not directly demonstrate regulation of *SICAvar* or *kir* by these pathways in *P. knowlesi*. Additional enriched terms included stress-associated categories such as response to heat, protein stabilization, and apicoplast-associated functions. Together with enrichment of protein-folding and organellar maintenance pathways, these patterns are consistent with activation of a stress-response–like transcriptional state (60). This interpretation is further supported by differential expression of HMGB2 and related chromatin-associated factors. Host body temperature at presentation was not associated with disease severity or parasite expression, nor was transcript expression directly associated with fever (Supplementary Table 14 and Supplementary Figure 12). By contrast, uncomplicated malaria cases did not exhibit an expansive stress-associated transcriptional activation. Instead, pathways linked to transcriptional and chromatin-level regulation were enriched, including chromatin binding, nuclear body organization, and promoter-specific control (Figure 3G), involving general transcriptional regulators such as histone methyltransferases, MORC, and SWI/SNF remodellers.

### Reciprocal genetic and transcriptional links underpin parasite disease phenotypes

To integrate parasite genetic and transcriptional variation, we performed eQTL analyses across samples with matched genomic and transcriptomic data (n = 96; Supplementary File 6). This bidirectional framework enabled us to link genetic associations identified by GWAS to regulatory effects on gene expression, while also identifying genetic variants underlying differentially expressed transcripts associated with disease severity and parasitaemia. We mapped *P. knowlesi* parasite eQTLs using FastLMM, applying both transcript-centric and SNP-centric models (Figure 4A). Cis-eQTLs were detected for several transcripts previously implicated by GWAS and differential expression, indicating that associated variants act, at least in part, through transcriptional regulation. For transcripts lacking confident mapping to the PkA1H1 reference genome, we employed a trans-eQTL framework, enabling inclusion of fragmented or highly polymorphic loci.

**Figure 4.**
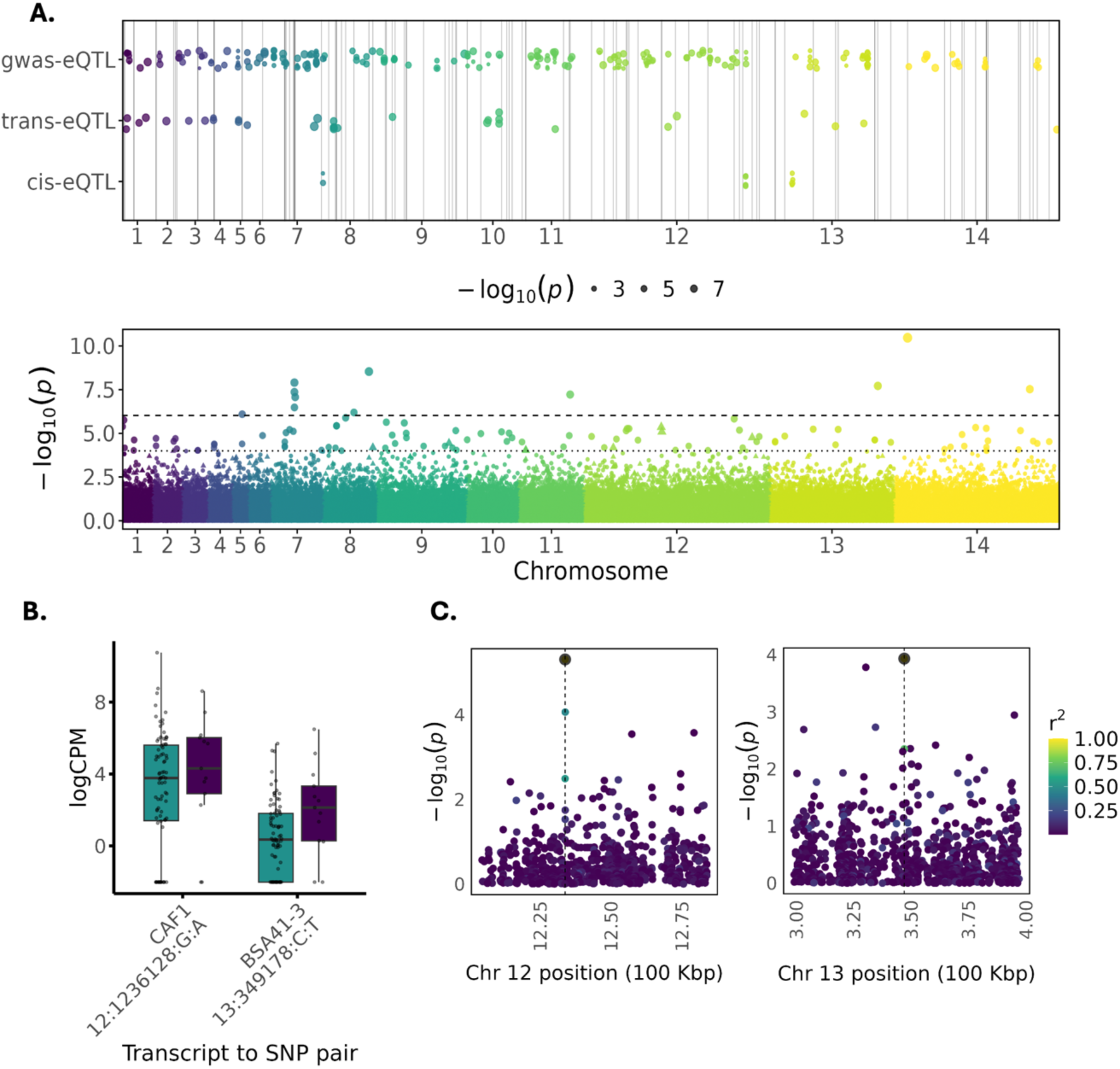
Integrative analysis of genome-wide association and transcriptomic data identifies regulatory loci in *Plasmodium knowlesi*. (A) Genome-wide distribution of eQTL signals identified using transcript- and SNP-centric models, stratified as cis-, trans-, or GWAS-linked eQTLs (top), shown alongside the Manhattan plot of combined severity and parasitaemia GWAS results (bottom). Horizontal lines indicate suggestive and genome-wide significance thresholds. **(B)** Expression levels (log-transformed counts per million) for two representative transcript–SNP pairs across uncomplicated and severe infections, illustrating phenotypic differences associated with regulatory variants. CAF1 denotes CCR4-associated factor 1; BSA41-3 denotes blood-stage antigen 41-3. **(C)** Regional association plots for the same loci shown in (B), displaying eQTL signal strength (−log₁₀ p) for variants within ±50 kb of the lead SNP. Points are coloured by linkage disequilibrium (LD; r²) with the lead variant, highlighting local genetic architecture underlying the regulatory signal.

Representative examples are shown in Figure 4B–C, including blood-stage antigen 41-3, which exhibited strong cis-associations with nearby SNPs, as well as regulatory variants identified for CCR4-associated factor 1. Beyond these illustrated loci, additional eQTL signals mapped to conserved and uncharacterised *Plasmodium* proteins, including loci on chromosomes 7 and 12. We detected significant eQTL signals involving a dynein heavy chain gene that was independently identified in the parasitaemia GWAS, linking genetic association at this locus to transcriptional variation. Regulatory effects involving members of the *SICAvar* multigene family were observed, connecting parasite regulatory variation to expression diversity within antigenically variable repertoires. Together, these analyses demonstrate convergence between parasite genetic variation and transcriptional programs associated with disease severity and parasitaemia, supporting a regulatory basis for key disease-linked pathways identified across GWAS and differential expression analyses.

## Discussion

Our multi-omic analysis identifies parasite-intrinsic programs associated with disease severity and parasite burden in clinical *Plasmodium knowlesi* infections. By combining GWAS, selection scans (XP-EHH), transcriptomics, and eQTL mapping, we link parasite genetic variation to coordinated transcriptional programs involving invasion, stress adaptation, immune evasion, and proliferation. In contrast to the long-studied human malaria parasites, *P. knowlesi* represents a rapidly emerging zoonotic pathogen whose transmission is sustained by macaque reservoirs and whose incidence continues to rise across Southeast Asia (1, 2, 3, 4, 5, 6). Understanding parasite factors that contribute to severe disease is therefore not only biologically important but also critical for anticipating the clinical impact and evolutionary trajectory of this expanding zoonosis. Our results suggest that parasite burden is primarily associated with invasion- and proliferation-linked programs, whereas severe disease is characterised by transcriptional signatures consistent with stress tolerance and immune evasion. Notably, these transcriptional patterns were independent of IDC stage composition and were partly genetically controlled through regulatory variation identified by eQTL mapping. Together, these findings indicate that clinical severity may reflect parasite-intrinsic regulatory states rather than parasite biomass or developmental timing alone, highlighting molecular pathways that could help track the emergence or spread of high-virulence parasite lineages in zoonotic malaria populations.

Importantly, this independence from IDC composition is consistent with the largely asynchronous biology of *P. knowlesi*. In contrast to *P. falciparum*, where sequestration of mature stages complicates interpretation of peripheral blood transcriptomes (61, 62), *P. knowlesi* permits direct observation of parasite-intrinsic regulatory programs in clinical samples. Deconvolution analysis confirmed trophozoites as the dominant life stage, with no systematic differences between severe and uncomplicated cases, reinforcing that IDC composition is not a major driver of severity. Mature male and female gametocytes were consistently detected, including in infections with very low parasitaemia, suggesting transmission potential in sub-microscopic or symptomatic cases of *P. knowlesi*. This raises the possibility of cryptic human reservoirs, which is particularly relevant in the context of *P. knowlesi*’s zoonotic biology: although sustained human-to-human transmission has not been observed in nature, it has been demonstrated experimentally in the laboratory (63, 64), suggesting that low-density infections may plausibly contribute to transmission under appropriate ecological and epidemiological conditions.

Severe *P. knowlesi* malaria is characterised by a coordinated transcriptional program consistent with stress adaptation and energetic trade-offs. Whereas parasitaemia-associated GWAS loci included helicases, cyclins, and peptidases–consistent with parasite-intrinsic determinants of proliferative capacity (39, 40, 41, 42)–severe infections were marked transcriptomically by upregulation of transcriptional, translational, and protein-folding machinery, alongside rescued expression of cell-cycle and apoptosis-linked pathways, suggesting a shift in cellular priorities. GO enrichment highlighted protein stabilization, vesicle-mediated transport, and apicoplast/mitochondrial maintenance, resembling the “adaptive heat-stress” phenotype described in *P. falciparum*, where wild-type parasites upregulate chaperones and organellar pathways to withstand febrile or immune pressure (60). These severity-associated signatures were independent of host temperature and life-stage composition, pointing to a parasite-intrinsic stress response. More broadly, this interpretation is consistent with the ecology of *P. knowlesi*, whose natural macaque hosts have a higher baseline body temperature than humans (65), suggesting that the severity-associated stress signature observed here is unlikely to reflect a direct response to host fever alone.

Genetic analyses reinforce this stress-adaptation model by implicating chromatin-associated regulators of transcriptional plasticity. The severity GWAS identified suggestive associations with a putative S-adenosylmethionine–dependent methyltransferase, implicating chromatin-mediated regulation under host pressure (38). Notably, the GWAS implicated CCR4-associated factor 1, while differential expression analysis highlighted a distinct CCR4–NOT complex subunit, suggesting convergence at the level of the CCR4–NOT regulatory complex, which regulates transcription and mRNA turnover and has been shown to be dispensable during prolonged growth in macaque erythrocytes (19), linking host-specific regulatory requirements to severe human infection. Transcriptomic upregulation of HMGB2, a chromatin protein and known alarmin, further suggests roles in transcriptional regulation and host inflammation (55). eQTL analysis additionally implicated regulatory variants for blood-stage antigen 41-3, a gene of undefined function but with orthology to *P. falciparum* and blood-stage expression, linking genetic variation to modulation of host-parasite interaction pathways (66). In contrast, uncomplicated *P. knowlesi* infections lacked the broad activation of translational and stress-associated pathways observed in severe disease. Together, these findings support a model in which severe infections reflect a transcriptionally expansive, stress-engaged parasite state under host pressure. Stress-response pathways may therefore represent candidate therapeutic vulnerabilities in infections exhibiting this expansive program, consistent with growing interest in targeting parasite stress-adaptation mechanisms in antimalarial drug development (67, 68, 69).

Parasite burden, rather than severity, is most strongly linked to transcriptional programs governing invasion and host–parasite interaction. While some parasites appear to prioritize stress adaptation, others maintain or amplify red blood cell invasive capacity, a feature most strongly associated with parasitaemia and only variably with severity. Genetic and transcriptomic analyses highlight multiple genes involved in host-parasite interactions and red blood cell invasion, suggesting that variation in invasion efficiency contributes to parasite burden and, indirectly, to clinical outcome. However, these transcripts likely reflect parasite-intrinsic growth potential or host cell remodelling, rather than severity, suggesting that some apparent severity associations are mediated through parasitaemia. Consistent with this interpretation, our findings align with those of *Moon et al.* and *Ahmed et al.*, as genes essential for human erythrocyte invasion are upregulated in severe infections, whereas genes associated with macaque invasion or slower-growing phenotypes in humans are preferentially expressed in uncomplicated cases (14, 70). Notably, the parasitaemia-associated DEAD-box helicase identified here has clear orthologues in human and other non-human primate malaria parasites but no detectable orthologue in rodent malaria species in current PlasmoDB annotations, suggesting potential lineage-specific differences in determinants of growth capacity. Together, these results support a continuum in which host-adapted invasion programs are reinforced during high-burden infections, even when underlying genetic associations are distributed across distinct loci. Efficient invasion promotes parasite expansion and increases the need for immune evasion to sustain high parasite densities and, potentially contributing to severe disease (71). We did not detect associations at canonical invasion complex components (e.g., PTRAMP/CSS/RIPR orthologues), suggesting that severe-disease risk is unlikely to be explained by discrete variation in these established invasion modules (72).

Beyond invasion ligands, genetic signals also implicate intracellular transport and cytoskeletal machinery in parasite expansion (73). Consistent with this, three dynein heavy chain (DHC) genes and a dynein light intermediate chain gene were associated with parasitaemia, implicating the dynein motor complex in sustaining high parasite burdens. In rodent malaria, DHC3 (orthologue of the PKA1H_070007900 GWAS hit) forms part of a subpellicular microtubule–associated dynein complex essential for apical cargo transport, morphogenesis, and gliding motility, with disruption blocking ookinete development and transmission (74). Although DHCs are not expressed in asexual blood stages of *P. yoelii* or *P. falciparum* (74, 75), dynein complexes localise to the apical pole of *P. falciparum* merozoites during late schizogony (76), and pharmacological dynein inhibition reduces invasion efficiency, implicating roles in late-stage maturation or apical trafficking. The essential dynein light chain PfDLC1 further supports a conserved requirement for this pathway in erythrocytic development (75).

Importantly, the detection of eQTL signals at a DHC locus provides additional evidence that genetic variation in this motor-complex pathway is coupled to altered transcriptional regulation, linking parasite genotype to molecular phenotype. While dynein complexes have established roles in cytoskeletal organisation, apical cargo transport, and invasion-related processes in other *Plasmodium* species (74, 75, 76), lineage-specific dynein variation has not previously been linked to *in vivo* differences in parasite growth capacity. Selection signals enriched among uncomplicated infections suggest that dynein-associated variation contributes to differential growth or regulatory states across clinical phenotypes. Together, GWAS, regulatory, and selection signals supports a model in which DHC variation modulates intracellular transport or late schizont maturation, thereby shaping parasite growth capacity. Rather than directly determining disease severity, such variation may constrain an upper limit of achievable parasite biomass, creating conditions under which immune-evasion programs, including *SICAvar* and *kir* activation, become necessary for persistence. These convergent genomic and transcriptomic signals define a tractable framework for experimental follow-up, in which regulatory and coding variation at dynein loci can be tested *in vitro* to determine their effects on transcription, intracellular transport, and parasite growth.

Downstream of these growth- and invasion-associated processes, parasitaemia was associated with upregulation of 13 histone-related transcripts, including a H3 ortholog (PKA1H_110038200; PfH3.3, PF3D7_0617900) and its canonical counterpart (PF3D7_0610400), suggesting chromatin-mediated regulation as a potential mechanism underlying repertoire activation. In *P. falciparum*, histone H3 acetylation is associated with active *var* transcription (77, 78), suggesting related epigenetic processes may operate in *P. knowlesi*. Consistent with this interpretation, variant gene family activation emerged across multiple analyses, implicating both the level and breadth of *SICAvar* and *kir* expression in parasite persistence and immune evasion. The *SICAvar* multigene family encodes PfEMP1-like variant surface antigens that, unlike the subtelomerically clustered *var* genes of *P. falciparum*, are dispersed across chromosomes in gene-sparse regions permissive to rapid evolution and diversification (79). In macaque infection models, *SICA* expression is lost following splenectomy, consistent with strong host-dependent regulation (80). Multiple *SICAvar* and *kir* transcripts were associated with parasite load, indicating that broader activation of variant antigen repertoires accompanies sustained parasitaemia, with severe disease representing one potential outcome along this continuum. The absence of strong locus-specific GWAS signals at individual SICAvar genes likely reflects their multicopy architecture, extreme sequence diversity, and coordinated epigenetic regulation at the family level rather than discrete single-locus effects.

Importantly, activation extended beyond increased expression levels to differences in repertoire breadth. While both *SICAvar* and *kir* families showed higher overall expression in severe and high-parasitaemia infections, only *kir* exhibited a significantly greater number of expressed transcripts per infection, indicating differences in activation dynamics rather than simple upregulation alone. Across *Plasmodium* species, effective immunity constrains variant antigen repertoire breadth, whereas reduced immune control permits broader activation. Consistent with this, macaque infection models show reduced *SICAvar* repertoire diversity under immune control (23, 81), while asymptomatic *P. falciparum* infections exhibit more focused *var* expression than clinical disease (82). Experimental infections with *P. chabaudi* further demonstrate that vector transmission resets expression of *pir* variant families—functional analogues of antigenic variation systems in other *Plasmodium* species—followed by gradual expansion of expressed variants during blood-stage infection before immune pressure narrows expression to a smaller subset (83). A similar process may underlie the broader *kir* repertoires observed here, potentially reflecting prolonged or poorly controlled infections. In contrast, the more heterogenous *SICAvar* signal across infections may reflect both biological constraints and technical factors, including greater sequence diversity and assembly fragmentation, which reduce power to detect transcript-level effects despite clear family-level activation. Together, these observations support a model in which variant antigen families are regulated through both genetic and transcriptional mechanisms, enabling parasite persistence and biomass accumulation, with severe disease emerging under conditions of reduced immune control.

Several limitations should be considered when interpreting these integrative genomic findings. First, all samples were derived from clinical infections in Sabah, Malaysian Borneo, and it remains unclear to what extent the findings generalise to other regions or transmission settings. Second, determining whether host-intrinsic factors influence parasite gene expression will require integrated host–parasite genomic and transcriptomic analyses. Third, parasite load was quantified using peripheral blood parasitaemia, which may not fully capture total parasite biomass, although evidence for non-circulating niches in *P. knowlesi* remains limited. Fourth, inference of life-cycle composition relied on cross-species reference-based deconvolution, which may not fully capture species-specific transcriptional differences. Finally, while consistent genetic signals were implicated across analyses, functional validation will be required to establish causal mechanisms. Despite these constraints, the convergence of genetic, transcriptional, and evolutionary signals highlights parasite regulatory variation as a tractable target for translational applications, including genomic surveillance for high-risk lineages, molecular risk stratification, and identification of parasite pathways that may influence interactions with host immunity or antimalarial interventions.

Our data converge on a model in which parasite regulatory variation, rather than parasite load alone, shapes disease phenotype in *P. knowlesi*. Parasitaemia was linked to upregulation of replication and invasion programs, including erythrocyte membrane–associated proteins and blood-stage antigens, whereas severity was marked by transcriptional signatures consistent with stress tolerance and immune evasion. Both variant antigen expression (*SICAvar* and *kir*) and invasion were more strongly associated with parasitaemia than severity, suggesting that immune evasion programs enable parasite persistence and biomass accumulation, with severe disease reflecting genetically encoded differences in regulatory response that manifest under particular host or ecological contexts. By linking transcriptional phenotypes to parasite genotype, eQTL mapping identifies regulatory variation as a central mechanism modulating growth, stress adaptation, and evasion *in vivo*, with results representing testable candidates in future mechanistic work. These findings advance understanding of *P. knowlesi* pathogenesis while highlighting regulatory loci and transcriptional programs that may inform genomic surveillance, risk stratification, and the monitoring of emergent high-risk parasite lineages. Beyond *P. knowlesi*, this integrative framework provides a generalizable strategy for dissecting parasite-intrinsic determinants of clinical outcome in non-*P. falciparum* and zoonotic malaria species, where comparable studies remain scarce.

## Methods

### Sample collection and genomic sequencing

We utilised a combination of newly generated *P. knowlesi* whole-genome sequencing data (84) and archived FASTQ files from samples of *P. knowlesi*-infected patients in Malaysia. Newly processed samples were collected as part of prospective clinical studies conducted through the Infectious Diseases Society Kota Kinabalu Sabah-Menzies School of Health Research collaboration from 2011 to 2018 across multiple hospital sites in Sabah (11, 12, 85). Patients of all ages presenting with microscopy-diagnosed malaria were enrolled following informed consent. *P. knowlesi* infections were confirmed through validated PCR (86, 87, 88) and parasitaemia quantified by expert research microscopists using the patient’s own white blood cell count from hospital haematology results (89). Severe malaria phenotype was defined for each participant using the WHO research criteria for *P. knowlesi* (34). 737 clinical isolates underwent Illumina whole genome, paired-end sequencing (150bp), with library preparation conducted using the NEBNext® Ultra™ IIDNA Library Prep Kit (from New England BioLabs Inc., Cat No. E7645). Ethical approval was obtained from the medical research ethics committees of the Ministry of Health, Malaysia and Menzies School of Health Research, Australia.

### Total RNA extraction and measurements

200 µl of packed red blood cells (pRBC) depleted of white blood cells from 402 field specimens were homogenized in 400 µl DNA/RNA Shield (ZYMO). 600 µl of TRIzol reagent (Invitrogen) and 120 µl of chloroform were added and tubes were centrifuged for 5 min at 20000 x g. Clear supernatant was aspirated and transferred to a new 2 ml 96-deep well plate (Corning). RNA was extracted using ZYMO DirectZol-96 MagBead RNA kit (ZYMO) following manufacturer instructions. RNA was eluted in 17 µl of RNAse/DNAse free water, and its purity was assessed by spectrometry on Nanodrop (ThermoScientific). The RNA concentration was estimated using RNA-specific Qubit fluorometric assays (Invitrogen). RNA integrity was assessed on Bioanalyzer RNA Nano Chip (Agilent). RNA quality cut-off metrics were as follow: 260/280 ratio ≥1.5, 260/230 ratio ≥1.5, RIN ≥4 (RIN_av_ ∼ 7), RNA yield ≥40ng. RNA samples that passed quality metrics were selected for sequencing. All steps were automated on Hamilton STAR liquid handling platform (Hamilton Robotics).

### Stranded total RNA library preparation

Prior to library preparation 40 ng of total RNA per each sample was DNase treated with TURBO DNase (ThermoFisher). Subsequently RNA libraries were prepared using ZYMO-Seq RiboFree Total RNA Library Kit (ZYMO) following the manufacturer’s instructions with some modifications to the protocol described below. This protocol generated stranded, rRNA, and human globin-depleted libraries. The depletion step of parasite rRNA and human globin was adjusted to 90 minutes at 68°C. 12 cycles of PCR were used for indexing. The resulting libraries were eluted in 15ul of the elution buffer and verified by real-time PCR, fluorometric DNA concentration assay (Invitrogen), and Bioanalyzer DNA High-Sensitivity chip (Agilent Technologies). 48 libraries were multiplexed, equimolarly pooled and sequenced on one lane of Novaseq X (Illumina) generating approximately 375GB of data per lane (Novogene Co., Singapore). Technical replicates of 3D7 *P. falciparum* strain RNA (rings, approx. 14hpi, 1% parasitaemia) were also included on each sequencing lane to control for library preparation and sequencing efficiency.

### Genomic variant calling and genome wide association analysis

Genotypes were generated as previously described (16) using a pipeline based on a previously published workflow (90) and following many of the Genome Analysis Toolkit’s (GATK) best practices (91). In brief, sequencing reads were aligned to the *P. knowlesi* PKA1-H.1 reference genome (17) using the Burrows-Wheeler Aligner (bwa). BAM pre-processing was performed with Picard version 2.26.1 and GATK version 3.8-1-0 (91). SNPs and indels were identified using a consensus approach, combining calls from GATK and bcftools with a modified version of a previously described workflow (92, 93). For GATK, HaplotypeCaller was used to call variants in each sample, followed by joint genotyping with CombineGVCFs and GenotypeGVCFs. A parallel joint-calling approach was implemented using bcftools mpileup and call. A consensus of the two call sets was taken to generate a conservative list of high-quality variants. SNPs and indels were filtered with GATK’s VariantFiltration using thresholds of FS ≤ 2, MQ ≥ 59, QD ≥ 20, QUAL ≥ 30, and depth ≥ 5, based on the distribution of these metrics across the dataset.

Genome-wide association analyses (GWAS) were performed separately for parasitaemia and severity phenotypes, using multiple SNP and sample sets. Model performance was evaluated using genomic inflation factors (λ). For all datasets, moimix (github.com/bahlolab/moimix) was used to calculate the within-isolate fixation index (Fws), and complex infections (Fws < 0.95) were excluded. Genotypic missingness, sample missingness, and minor allele frequency were calculated in PLINK (94, 95), which was also used to apply a range of filtering thresholds, from stringent (genotypic & sample missingness < 5%) to relaxed (< 20%). Warped fastLMM (96) was applied for association testing on each dataset. For the best-performing models (λ closest to 1), genome-wide significance thresholds were defined using a Bonferroni correction based on the LD-pruned SNP set used to construct the fastLMM distance matrix. LD pruning was performed with a 150 bp window, shifting by 5 SNPs and a variance inflation factor (VIF) of 2 in PLINK. A more relaxed suggestive threshold (*p* < 1x10^-4^) was also defined to capture high-ranking loci, justified on the basis of the *P. knowlesi* genome size (∼23 Mb) and the corresponding approximate number of independent association tests. Gene essentiality was inferred from a published supersaturated piggyBac transposon mutagenesis screen in *P. knowlesi* A1H1 (36), where genes lacking any recovered CDS insertions were classified as essential.

For selected *P. knowlesi* proteins with unknown function but strong GWAS association signals, full-length amino acid sequences were retrieved from PlasmoDB (35). Protein structure prediction was performed using AlphaFold2 via the ColabFold notebook (v2.3.0) (97), using default settings and MMseqs2-based MSA generation. Five models were generated per sequence, and model confidence was assessed via per-residue pLDDT scores and predicted aligned error (PAE). Structural homology searches were conducted using FoldSeek (98), with default search parameters against all available databases. Structural matches with high FoldSeek probability (>0.9) and overlapping high-confidence AlphaFold regions (pLDDT > 90) were retained for interpretation.

### *De novo* transcriptomic assembly and transcript abundances

For transcriptomic analyses, we constructed a *de novo* meta-transcriptome assembly using isolates collected in this study. Although a high-quality reference genome exists for *P. knowlesi*, transcriptome annotations remain incomplete, particularly for hypervariable multigene families. A *de novo* strategy was therefore adopted to minimize reference bias, recover poorly annotated gene families such as *SICAvar*, and capture novel or sample-specific transcripts and isoforms relevant to host adaptation and immune evasion. Transcriptome assembly was performed using a Trinity-based workflow implemented within a Snakemake pipeline. Standard quality control and adapter removal were carried out with FastQC and BBDuk, followed by alignment of cleaned reads to both the human genome (to remove host contamination) and the *P. knowlesi* reference genome (for additional quality assessment) using the STAR aligner (99). The remaining reads were assembled with Trinity (47), and redundancy was reduced using CD-HIT-EST (100, 101, 102). Transcript abundance estimation and count matrix generation were performed using the Trinity utility scripts *align_and_estimate_abundance.pl* and *abundance_to_matrix.pl*. Assembly quality was evaluated using Trinity’s built-in metrics.

### Assembly annotation

Following assembly, transcripts were functionally annotated using a hybrid strategy that combined reference-based mapping to the *P. knowlesi* PKA1H1 genome with *de novo* annotation through the Trinotate framework, enabling comprehensive assignment of coding, non-coding, and structural features. Annotation was performed using Trinotate (49), a comprehensive framework for Trinity assemblies that integrates multiple predictive tools to generate functional annotations. TransDecoder (103) was first used to identify putative open reading frames, and an SQLite (104) database was built to store predicted coding regions together with the *de novo* assembly and assembly map. Transcript sequences were searched against the UniProt/Swiss-Prot database (105) using DIAMOND (106) to identify homologies to known proteins, while conserved protein domains were detected with HMMER (107) against the Pfam-A database. Additional functional predictions included SignalP (108) for signal peptides, TMHMM (109) for transmembrane helices, and RNAMMER (110) for ribosomal RNA features. To capture structured RNA elements and potential intergenic transcripts, we further applied Infernal (111) with the Rfam database, enabling identification of conserved RNA motifs beyond coding regions. All results were integrated into the SQLite database by Trinotate, which generated the final annotation report, including Gene Ontology (GO) terms (112, 113), KEGG pathway assignments (114, 115), and other relevant functional annotations.

To assign a single representative annotation to each transcript, we combined results from reference-based and *de novo* annotation workflows and applied a hierarchical prioritization scheme. Where available, annotations from the *P. knowlesi* PKA1H1 reference genome were given highest priority. For transcripts without PKA1H1 annotation, we sequentially selected the best-supported match from Trinotate outputs, prioritizing UniProt/Swiss-Prot BLASTX hits, then BLASTP hits, followed by Pfam domain predictions, SignalP, TMHMM, and KEGG assignments. Transcripts lacking protein-based annotation were further queried against Rfam models using Infernal, enabling assignment of conserved structural RNA features. This procedure ensured each transcript was represented by a single concise annotation in downstream analyses, while full annotation tables, including all supporting evidence, are provided in the Supplementary Information.

### Intra erythrocytic cycle deconvolution

We estimated IDC stage composition in bulk *P. knowlesi* RNA-seq samples using Scaden, a deep learning–based cell-type deconvolution framework (50). Single-cell references were derived from the Malaria Cell Atlas datasets for *P. knowlesi* (10x Chromium dataset spanning ring, trophozoite, schizont), *P. falciparum* and *P. berghei* (10x Chromium dataset spanning ring, trophozoite, schizont, and male, female, and developing gametocytes) (21, 22, 116). To enable cross-species comparisons, we generated a one-to-one ortholog map between genomes and restricted both bulk and single-cell matrices to shared orthologous genes. Bulk RNA-seq reads were aligned with STAR, and gene counts were converted to transcripts per million (TPM) without log transformation to match Scaden input requirements. Reference single-cell matrices were processed as AnnData objects (.h5ad) with stage labels, then filtered to the ortholog gene set. For each reference dataset, we performed 10 independent Scaden runs to stabilize estimates. In each run, 1,000 simulated pseudo-bulk mixtures were generated from the single-cell reference, used jointly with the bulk data for training (5,000 steps), and then applied to predict stage proportions in each bulk sample. Predictions were averaged across runs to obtain final stage estimates, and the run-to-run variance was used as a measure of deconvolution stability. Comparisons between severe and uncomplicated malaria groups were performed using Wilcoxon rank-sum tests with Benjamini–Hochberg correction, and associations with parasitaemia were assessed using Spearman correlation.

### Differential expression analysis

Differential expression analysis was performed using a consensus approach adapted from consensusDE (92), combining results from DESeq2 (117), edgeR (118), and voom-limma (119) to identify robust differentially expressed transcripts. Trinity outputs were converted into HTSeq-compatible format using custom Python scripts, and a SummarizedExperiment object was generated through consensusDE. Initial filtering was applied with buildSummarized(filter = TRUE), which uses edgeR’s CPM-based method to remove lowly expressed transcripts (CPM >1 in at least 10 samples). Samples without group annotations and transcripts with zero counts were excluded, and highly expressed outliers (mean counts > 50,000) were removed to avoid skewing results. To control for unwanted variation, we applied the RUVr method from RUVSeq (120): a first-pass GLM was fitted using sample groupings and sequencing batch as a random effect, and deviance residuals were used to estimate one factor of unwanted variation (*k* = 1), which was then added to the sample metadata. Transcripts with missing values were excluded, and all filtering steps were logged for reproducibility.

Differential expression was carried out with DESeq2, edgeR, and voom-limma, each incorporating RUV factors and sequencing batch as independent effects. DESeq2 used shrinkage-based dispersion estimation with generalized linear modeling, edgeR employed quasi-likelihood fitting, and voom applied precision weighting of log-transformed counts prior to linear modelling. Diagnostic plots (p-value histograms, MA plots, dispersion estimates, volcano plots) were generated for each method to assess model behaviour. Principal component analysis (PCA) on normalized data was used to evaluate sample clustering and detect batch effects or outliers, with additional PCA performed on significant genes and the top 1,000 most variable genes. Heatmaps of top-ranked DEGs genes provided visualization of group-level differences. To ensure biological relevance, we further filtered the consensus set of significant transcripts (adj-*p* < 0.05) to retain only those expressed in at least 30% of samples in either group. Concordance across methods was assessed by overlap analysis of significant transcripts.

We manually compared differentially expressed genes with top GWAS hits and suggestive loci for severe malaria. Overlap was defined based on gene name similarity, protein domain conservation, or known functional alignment (e.g., invasion, egress, transcriptional regulation). This approach was used to highlight potential biological convergence rather than direct genomic co-localisation.

### Targeted exploratory transcriptomic analysis

To investigate the expression dynamics of specific gene families and functional groups, we performed targeted exploratory analyses using curated transcript subsets. Gene lists of interest (e.g., *SICAvar*, *kir*, host-related, and stress-response genes) were compiled from literature and functional annotations and matched to Trinity transcript IDs in the expression matrix. Analyses were performed using the pre-processed and filtered expression object generated for differential expression, as well as additional filtering for transcript prevenance (>30% of samples). For each gene set, we visualized expression using heatmaps of z-score-transformed logCPM values, calculated on a per-gene basis. In sample-level heatmaps, both transcripts and samples were clustered using hierarchical clustering of z-scores. In grouped heatmaps, genes were ordered by average expression in the severe malaria group.

### Gene Ontology (GO) enrichment analysis

To identify biological processes, molecular functions, and cellular components associated with gene expression changes, we performed Gene Ontology (GO) enrichment analysis using the gseGO function from the *clusterProfiler* R package (121, 122). This approach applies gene set enrichment analysis to a ranked list of transcripts, avoiding the need for an arbitrary differential expression cutoff and providing sensitivity to subtle, coordinated expression shifts. GO annotations were compiled from multiple sources, including *P. knowlesi* PKA1H1 reference gene models (PlasmoDB GAF file) and homology-based annotations derived from Trinotate (BLASTX, BLASTP, and Pfam). To reduce redundancy and prioritise curated sources, GO terms were assigned to each transcript based on a hierarchical preference: PKA1H1 > BLASTX > BLASTP > Pfam. The gene ranking metric was the signed –log₁₀(p-value), weighted by the direction and strength of differential expression (log₂ fold-change). Significantly enriched GO terms were defined using a false discovery rate (FDR) threshold of 0.01. Annotation source information was retained to aid interpretation of enriched terms. To visualise enrichment patterns, we generated GeneRatio dot plots summarising top-ranked terms, and heatmaps of mean z-scored expression for leading-edge transcripts across groups

### Expression quantitative trait loci (eQTL) analysis

We performed eQTL mapping using linear mixed models implemented in FastLMM, separately for each expressed transcript. Gene expression data were filtered using the same steps described for the differential expression analysis and quantified as log-transformed counts per million (log2-CPM). To avoid spurious associations driven by highly expressed housekeeping genes, transcripts above the 99th percentile of mean expression were excluded. Genotype data consisted of high-quality SNPs (genotypic missingness < 5%, sample missingness < 5%, and MAF > 5%) called from whole-genome sequencing of matched *P. knowlesi* isolates. To account for population structure, we included a precomputed LD-pruned genotype matrix as a random effect. The first five principal components from both the genotype and expression data were also included as fixed-effect covariates.

For transcripts with confident mapping to the *PkA1H1* reference, we performed targeted cis-eQTL scans by testing all SNPs genome-wide and then post-filtering to retain only those within a ±10 kb window of the transcript. P-values were Bonferroni-corrected per transcript using the number of cis-SNPs tested, with associations considered significant if corrected p < 0.05 and suggestive if nominal p < 1/N (where N is the number of cis-SNPs). Manhattan and QQ plots were generated only for transcripts with at least one suggestive hit. For differentially expressed transcripts that lacked unambiguous mapping to the *PkA1H1* reference, we conducted exploratory trans-eQTL scans. Here, each transcript was tested against all SNPs genome-wide, with associations evaluated after correction for multiple testing. This approach allowed us to capture potential long-range regulatory effects on genes that are not represented in the reference annotation but may still be biologically important.

Finally, to complement these transcript-centric scans, we also performed a SNP-centric analysis anchored on genome-wide association study (GWAS) hits. For each SNP identified as significant or suggestive in the GWAS, we queried local transcriptomic effects within a ±5–10 kb window using linear models within the FastLMM framework, again controlling for population structure and expression PCs. This enabled the identification of putative regulatory effects linked to top GWAS loci that may not be recovered in transcript-wide screens.

## Supporting information

Supplementary figures and tables supporting genetic, transcriptomic, and life-stage analyses.

Extended GWAS results for severity and parasitaemia, including summary statistics and selection analyses.

Differential expression results for severe vs uncomplicated malaria with functional annotations.

Differential expression results with parasitaemia as a continuous covariate and functional annotations.

Gene Ontology enrichment results and associated gene sets for pathway-level interpretation.

Integrative eQTL results linking parasite genetic variation to transcriptional regulation.

## Data Availability

Genomic data produced as part of this work are available at the Sequence Read archive (SRA) of the National Center for Biotechnology Information (NCBI) under the BioProject ID PRJNA1066389 and all bioinformatic and analytical scripts are available at https://github.com/JacobAFW/knowlesi_integrative_genomics#.

## Acknowledgements

We thank the study participants, and the research team at the Infectious Disease Society Kota Kinabalu Sabah including Sitti Saimah binti Sakam, Azielia Elastiqah binti Salamth and Mohd Rizan Osman. We thank the Director-General, Ministry of Health, Malaysia, for permission to publish this manuscript.

## Contributions

MJG, MAF, CD, NMA and SA conceived the study. MJG, MAF, ZB, SA, BEB, CD, GSR, NMA and TW acquired the project funding and contributed to study management. MJG, NMA, BEB, AMM, AJ, CD, TM and GSR coordinated clinical sample collection. MJG, MAF, ZB, SA, EDB, RA, SN, MK and JAFW conceptualised and designed the methodology. KP, MK, DTQH, ZB and JAFW coordinated or performed sample processing and sequencing library preparation. SN, MK, DA and JAFW curated the data. JAFW developed all computational pipelines and analysis scripts, and performed all data analyses, with intellectual input from EDB, MAF, SN, MK, HT, KSH, RM, ZB and MJG. JAFW drafted the manuscript, which was reviewed by all authors.

## Ethics Approval Statement

The research was conducted in accordance with the Declaration of Helsinki and ethics approval was obtained from the medical research ethics committees of the Ministry of Health, Malaysia and the Menzies School of Health Research, Australia.

## Supplementary Information

Supplementary File 1.

Supplementary figures and tables referenced in the main text, including additional genetic, transcriptomic, life-stage deconvolution, Alpha-Fold predictions, and quality-control analyses supporting the primary results.

Supplementary File 2.

Extended genome-wide association study (GWAS) results for clinical severity and parasitaemia, including full summary statistics for all tested variants, results of XP-EHH selection scans, exploratory analyses of top-ranked loci, and functional annotations based on transposon insertion essentiality data from PlasmoDB.

Supplementary File 3.

Extended results of differential expression analysis comparing severe and uncomplicated malaria, including consensus results and *P. knowlesi* reference and Trinotate functional annotations.

Supplementary File 4.

Extended results of differential expression analysis treating parasitaemia as a continuous covariate, including consensus results and *P. knowlesi* reference and Trinotate functional annotations.

Supplementary File 5.

Extended Gene Ontology (GO) enrichment results, including the top 50 enriched terms per molecular category and corresponding gene sets used for pathway-level interpretation.

Supplementary File 6.

Extended results of integrative eQTL analyses, including cis- and trans-eQTLs identified using both transcript-centric and SNP-centric models, linking parasite genetic variation to transcriptional regulation across severity- and parasitaemia -associated loci.

