## Supplementary figures and tables supporting genetic, transcriptomic, and life-stage analyses. for "Integrative genomics of *Plasmodium knowlesi* reveals parasite-intrinsic regulators of severe human malaria"

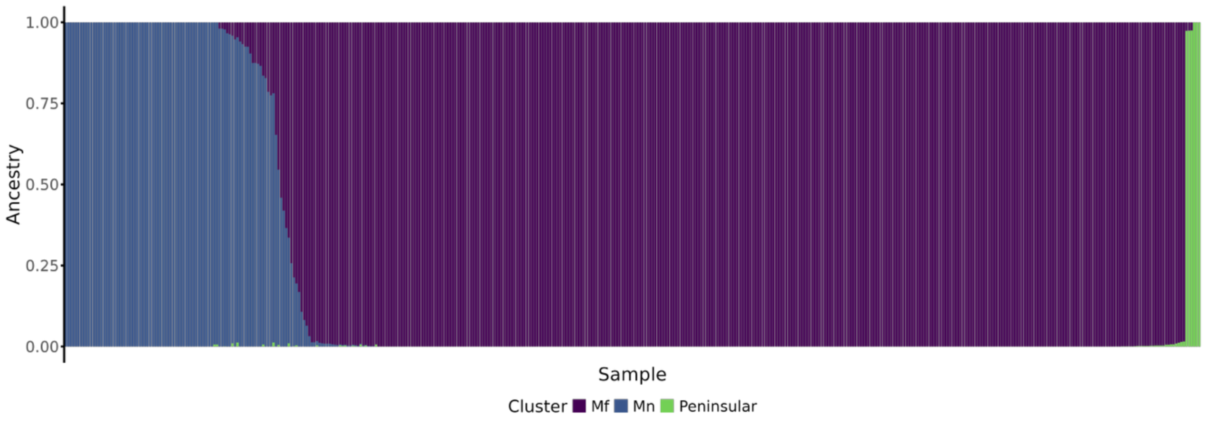


**Supplementary Figure 1. Inferred population structure of *P. knowlesi* isolates based on ADMIXTURE clustering.** Each vertical bar represents an individual isolate, with colours indicating proportional ancestry from inferred ancestral populations. Clustering highlights distinct genetic substructure consistent with geography and/or host origin.

| Group | n | Male, n (%) | Female, n (%) | Age (years), median (IQR) | Parasitemia, median (IQR) |
|---|---|---|---|---|---|
| All | 425 | 334 (78.6) | 90 (21.2) | 41.0 (26.0-51.0) | 9.07e+03 (4.36e+03-2.64e+04) |
| Severe | 112 | 92 (82.1) | 20 (17.9) | 47.5 (36.8-60.0) | 4.74e+04 (1.65e+04-1.44e+05) |
| Uncomplicated | 313 | 242 (77.3) | 70 (22.4) | 37.0 (24.0-49.0) | 6.59e+03 (3.79e+03-1.38e+04) |

**Supplementary Table 1. Clinical and demographic characteristics of participants included in genetic analyses of *Plasmodium knowlesi* infection.** Severe malaria was defined according to WHO criteria for *P. knowlesi* (Methods). Values are presented as n (%) or median (IQR).

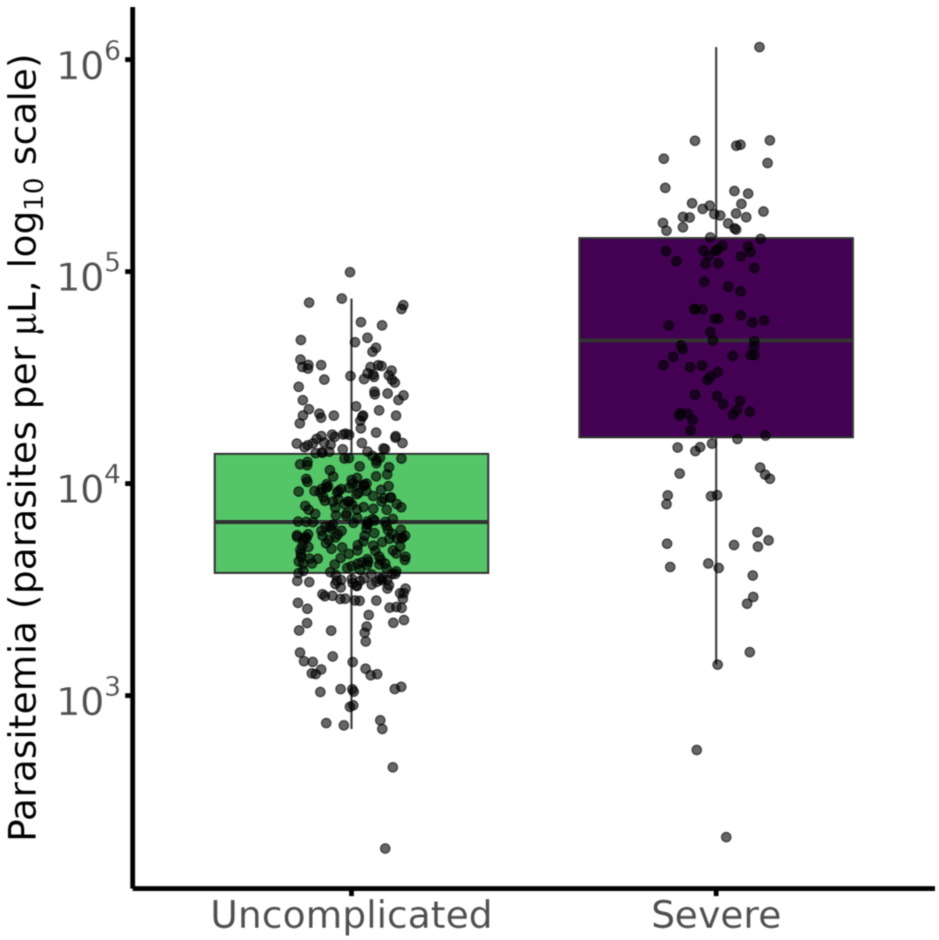


**Supplementary Figure 2. Parasitemia differs between severe and uncomplicated Plasmodium knowlesi infections.** Parasitemia (parasites per µL) is shown for patients with severe and uncomplicated malaria. Points represent individual patients and boxes indicate the median and interquartile range. Parasitemia was significantly higher in severe infections compared with uncomplicated infections (Wilcoxon rank-sum test, $P < 0.0001$), although substantial overlap between groups was observed.

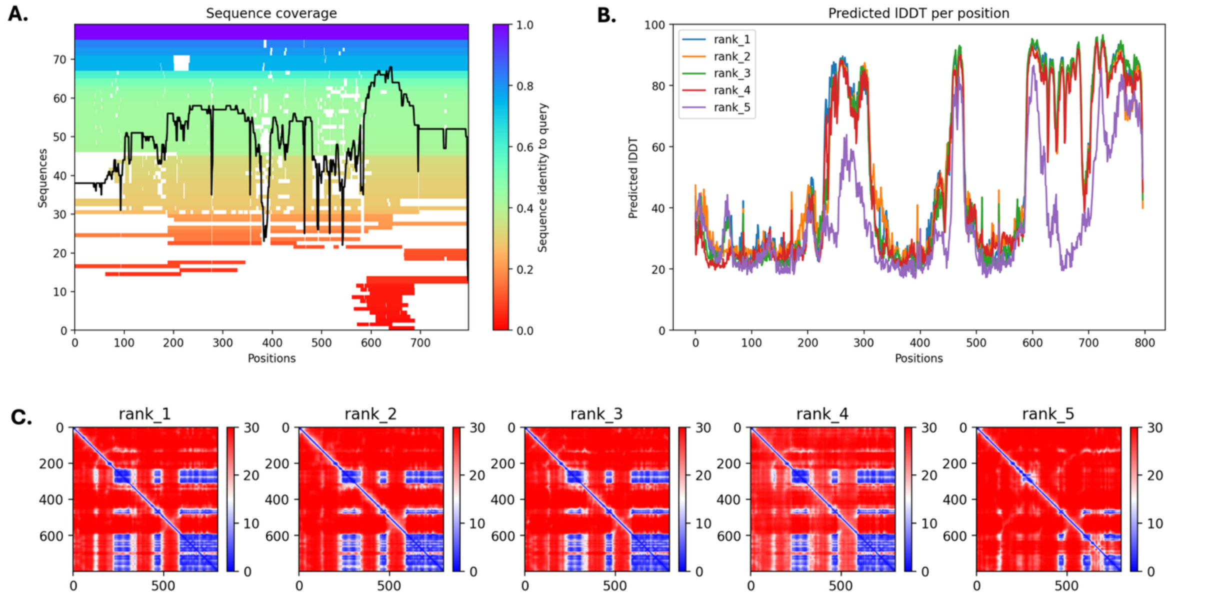


**Supplementary Figure 3. Structural prediction and confidence metrics for uncharacterized P. knowlesi protein PKA1H_070012000.** (**A**) Multiple sequence alignment (MSA) coverage and sequence identity for each aligned homolog used in structure prediction, with black line indicating number of aligned sequences per residue. (**B**) Per-residue predicted confidence scores (pLDDT) across five AlphaFold2 models, highlighting multiple high-confidence folded domains (pLDDT > 90). (**C**) Predicted Aligned Error (PAE) matrices for all five ranked models, indicating domain-level independence and potential modular structure.

| Target | Scientific Name | Prob. | Seq. Id. | E-Value | Score | Query Pos. |
|---|---|---|---|---|---|---|
| A0A481Z9L5 | Pithovirus LCPAC304 | 1 | 6.2 | 1.94E-06 | 210 | 591-764 |
| A0A7D3UQY4 | Fadolivirus algeromassiliense | 1 | 12.5 | 1.84E-06 | 198 | 564-760 |
| A0A2H4UVA7 | Bodo saltans virus | 1 | 11.2 | 7.30E-05 | 174 | 593-764 |
| A0A3G4ZUJ5 | Barrevirus sp. | 1 | 9.6 | 6.22E-05 | 168 | 591-761 |
| A0A3G5ADB4 | Satyrvirus sp. | 1 | 13.2 | 1.84E-06 | 159 | 572-762 |
| M1IM71 | Acanthocystis turfacea Chlorella virus OR0704.3 | 1 | 10.9 | 8.06E-04 | 123 | 593-762 |
| A0A0P0YMZ6 | Yellowstone lake phycodnavirus 2 | 0.99 | 11.3 | 2.11E-03 | 85 | 593-788 |
| A0A8T9VR71 | Iridoviridae | 0.97 | 11 | 5.22E-03 | 75 | 599-791 |

**Supplementary Table 2. High-confidence structural matches for PKA1H_070012000 identified by FoldSeek using AlphaFold2-predicted structures.** Structural matches were retained if FoldSeek probability >0.9 and overlapping regions had AlphaFold pLDDT >90. For each protein, model confidence, aligned region, structural hit, database source, and putative functional annotation are reported.

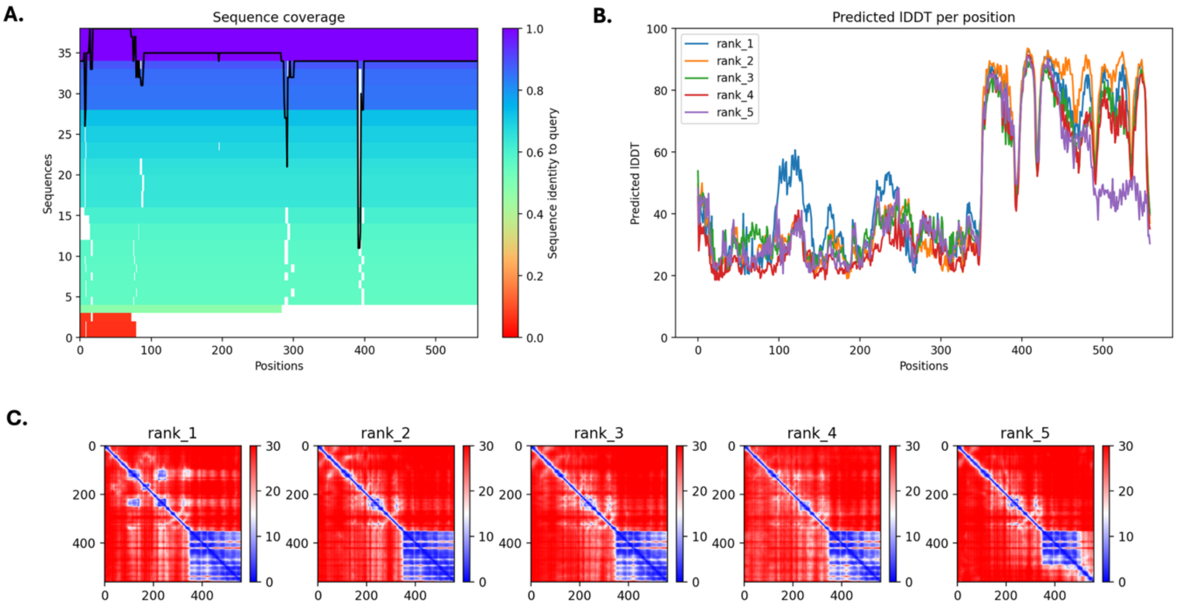


**Supplementary Figure 4. Structural prediction and confidence metrics for uncharacterized P. knowlesi protein PKA1H_070016000.** (**A**) Multiple sequence alignment (MSA) coverage and sequence identity for each aligned homolog used in structure prediction, with black line indicating number of aligned sequences per residue. (**B**) Per-residue predicted confidence scores (pLDDT) across five AlphaFold2 models, highlighting a single high-confidence folded domain (pLDDT > 90). (**C**) Predicted Aligned Error (PAE) matrices for all five ranked models, indicating domain-level independence and potential modular structure.

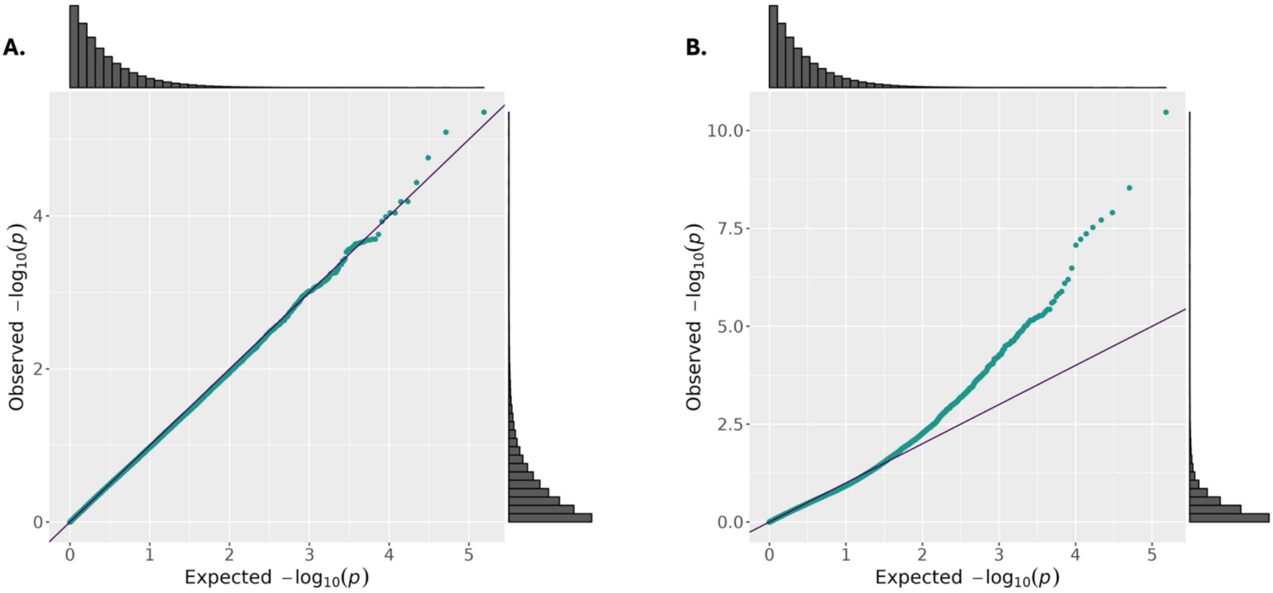

**Supplementary Figure 5. QQ plots of transcriptome-wide differential expression analyses for severity and parasitemia.** (A) Quantile–quantile (QQ) plot of observed versus expected p-values from the consensus differential expression model comparing severe versus uncomplicated malaria. The distribution largely follows the null, with deviation among the most significant transcripts consistent with biological signal ($\lambda = 0.998$). (B) QQ plot of p-values from models testing correlation between transcript expression and parasitemia. The upward deflection indicates widespread transcriptional association with parasite load, consistent with biological relevance of parasitemia-linked expression patterns ($\lambda = 0.960$). Marginal histograms show the distribution of observed $-\log_{10}(p)$ values in each analysis.

| Metric | Reference genome | De novo transcriptome |
|---|---|---|
| Median input reads | 7,209,880 | 7,209,880 |
| Input reads (Q25) | 4,778,271 | 4,778,271 |
| Input reads (Q75) | 20,729,931.75 | 20,729,931.75 |
| Median mapped reads | 1,957,322.5 | 4,031,267 |
| Mapped reads (Q25) | 352,252.5 | 1,623,161.25 |
| Mapped reads (Q75) | 8,844,753.5 | 16,024,962.25 |
| Median mapped (%) | 27.6 | 56.7 |
| Mapped % (Q25) | 8.7 | 34.1 |
| Mapped % (Q75) | 48.6 | 82.6 |

**Supplementary Table 3.** Comparison of read mapping rates to the P. knowlesi reference genome and de novo assembled transcriptome. Mapping statistics summarised across 210 clinical RNA-seq samples. Values represent the median and interquartile range (IQR; 25th–75th percentile) for input reads, mapped reads, and mapping percentages. Mapping to the de novo assembled transcriptome substantially increased read recovery relative to mapping to the reference genome.

| Dataset | Median library size (Q1–Q3) | Median genes detected (Q1–Q3) |
|---|---|---|
| Raw | 3,631,804 (1,379,695–16,137,502) | 66,612 (49,361–97,722) |
| Filtered | 543,392 (180,367–1,803,308) | 7,250 (6,207–7,938) |

**Supplementary Table 4.** Sequencing and transcript detection metrics before and after filtering. Median (Q1–Q3) per-sample values are shown for library size (total assigned reads), percentage mapped reads (reference and de novo assemblies), and number of detected transcripts (≥1 read), comparing the raw dataset and the final filtered analysis dataset.

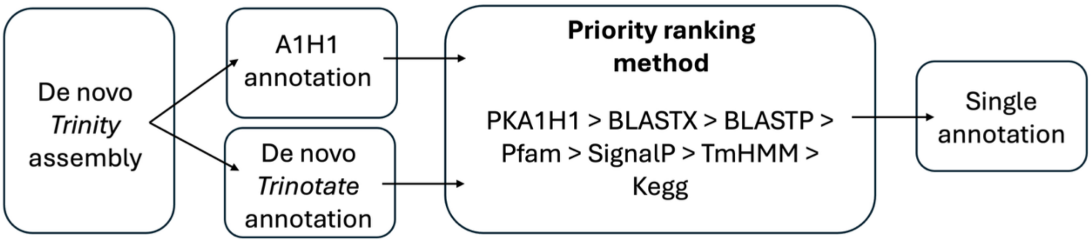


**Supplementary Figure 6. Annotation workflow showing integration of de novo *Trinity* assembly with PKA1H1 reference-guided and *Trinotate* annotation, prioritised by a structured ranking method (A1H1 > BLASTX > BLASTP > Pfam > SignalP > TmHMM > KEGG).**

| Life stage | W (on) | p-value (Wilcoxon) | Rho (Spearman) | p-value (Spearman) |
|---|---|---|---|---|
| Ring | 2951 | 0.26 | -0.12 | 0.08 |
| Schizont | 3121 | 0.53 | -0.03 | 0.64 |
| Trophozoite | 3433 | 0.77 | 0.06 | 0.35 |
| Gametocyte, developing | 3691 | 0.30 | -0.01 | 0.84 |
| Gametocyte, female | 3814 | 0.16 | 0.13 | 0.07 |
| Gametocyte, male | 3448 | 0.74 | -0.02 | 0.76 |

**Supplementary Table 5. Results for statistical testing of associations between Scaden-inferred IDC life stages and both severity and parasitemia.** *P. knowlesi* reference data was used for the asexual prediction and *P. berghei* for the sexual stages (1). Associations with severity were tested using Wilcoxon rank-sum tests and Spearman correlation analysis for $log_{10}$-transformed parasitemia.

| Life stage | W (Wilcoxon) | p-value (Wilcoxon) | Rho (Spearman) | p-value (Spearman) |
|---|---|---|---|---|
| Ring | 4175 | **0.01** | -0.02 | 0.80 |
| Schizont | 3535 | 0.56 | 0.05 | 0.45 |
| Trophozoite | 2639 | 0.04 | 0.01 | 0.92 |

| | | | | |
|---|---|---|---|---|
| Gametocyte, developing | 2696 | 0.06 | -0.01 | 0.91 |
| Gametocyte, female | 3240 | 0.78 | 0.14 | **0.05** |
| Gametocyte, male | 2894 | 0.20 | -0.01 | 0.90 |

**Supplementary Table 6. Results for statistical testing of associations between Scaden-inferred IDC life stages and both age and sex.** *P. knowlesi* reference data was used for the asexual prediction and *P. berghei* for the sexual stages (1). Associations with sex were tested using Wilcoxon rank-sum tests and Spearman correlation analysis for $\log_{10}$-transformed age.

| Parasitemia | Gametocyte, developing | Gametocyte, female | Gametocyte, male |
|---|---|---|---|
| 0 | 0.05 | 0.08 | 0.06 |
| 52 | 0.14 | 0.14 | 0.21 |
| 291 | 0.20 | 0.11 | 0.14 |
| 1040 | 0.03 | 0.04 | 0.03 |
| 1044 | 0.06 | 0.07 | 0.07 |
| 1175 | 0.09 | 0.08 | 0.13 |
| 1262 | 0.01 | 0.02 | 0.02 |
| 1274 | 0.03 | 0.05 | 0.03 |
| 1400 | 0.06 | 0.08 | 0.06 |
| 1440 | 0.03 | 0.05 | 0.04 |

**Supplementary Table 7. Low parasitemia (< 5000 parasites/µL) samples with proportions of gametocytes predicted using Scaden with the *P. berghei* reference data.** Parasitemia is reported as parasites per microliter.

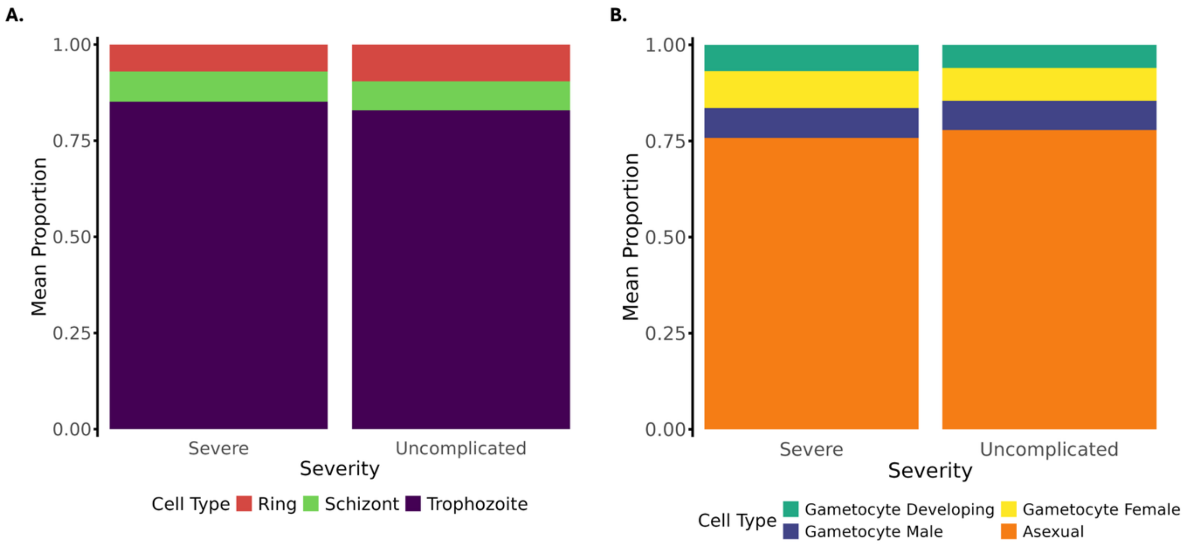


**Supplementary Figure 7. Mean Intraerythrocytic developmental cycle (IDC) composition of *Plasmodium knowlesi* infections.** Proportions of mean stage composition stratified by severe and uncomplicated infections, showing predicted gametocyte life stage proportions estimated with Scaden using *P. knowlesi* **(A)** and *P. berghei* **(B)** single cell reference sets.

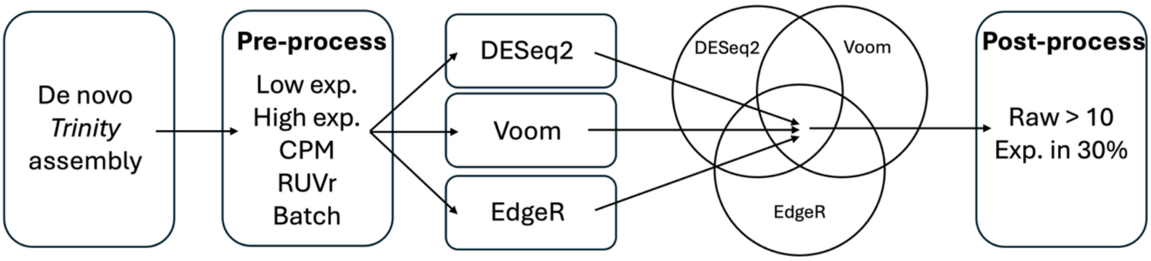


**Supplementary Figure 8. Differential expression (DE) analysis pipeline combining DESeq2, edgeR, and voom, with consensus outputs.**

| PC | Variable | $R^2$ (linear model) | p-value (linear model) | Rho (Spearman) | p-value (Spearman) |
|---|---|---|---|---|---|
| 1 | Library size | 0.011 | 0.127 | -0.097 | 0.161 |
| 2 | Library size | 0.013 | 0.093 | -0.166 | 0.016 |
| 3 | Library size | 0.002 | 0.531 | 0.027 | 0.7 |
| 1 | Transcripts detected | 0.092 | < 0.001 | 0.292 | < 0.001 |

| | | | | | |
|---|---|---|---|---|---|
| 2 | Transcripts detected | 0.05 | 0.001 | -0.231 | 0.001 |
| 3 | Transcripts detected | 0.49 | < 0.001 | 0.575 | < 0.001 |
| 1 | Mapped reads | 0.027 | 0.017 | 0.159 | 0.022 |
| 2 | Mapped reads | 0.054 | 0.001 | -0.228 | 0.001 |
| 3 | Mapped reads | 0.137 | < 0.001 | 0.181 | 0.009 |

**Supplementary Table 8. Correlation analyses comparing relationship between technical factors and principal components.** Results for several linear models and Spearman correlation analysis for library size, transcript counts and mapped reads vs the first three principal components.

| PC | Variable | $R^2$ (linear model) | p-value (linear model) | Rho (Spearman) | p-value (Spearman) |
|---|---|---|---|---|---|
| 1 | Schizont | 0.015 | 0.08 | -0.105 | 0.13 |
| 2 | Schizont | 0 | 0.976 | -0.026 | 0.704 |
| 3 | Schizont | 0.047 | 0.002 | -0.001 | 0.984 |
| 1 | Ring | 0.083 | < 0.001 | -0.437 | < 0.001 |
| 2 | Ring | 0.05 | 0.001 | -0.124 | 0.073 |
| 3 | Ring | 0 | 0.934 | 0.025 | 0.718 |
| 1 | Trophozoite | 0.089 | < 0.001 | 0.257 | < 0.001 |
| 2 | Trophozoite | 0.036 | 0.006 | -0.084 | 0.228 |
| 3 | Trophozoite | 0.01 | 0.153 | -0.055 | 0.426 |
| 1 | Developing gametocyte | 0.063 | < 0.001 | 0.318 | < 0.001 |
| 2 | Developing gametocyte | 0.063 | < 0.001 | 0.271 | < 0.001 |
| 3 | Developing gametocyte | 0.058 | < 0.001 | -0.084 | 0.225 |
| 1 | Female gametocyte | 0.038 | 0.005 | 0.251 | < 0.001 |
| 2 | Female gametocyte | 0.001 | 0.639 | 0.139 | 0.044 |
| 3 | Female gametocyte | 0.017 | 0.062 | -0.064 | 0.355 |
| 1 | Male gametocyte | 0.042 | 0.003 | 0.328 | < 0.001 |
| 2 | Male gametocyte | 0.026 | 0.02 | 0.256 | < 0.001 |
| 3 | Male gametocyte | 0.121 | < 0.001 | -0.186 | 0.007 |

**Supplementary Table 9. Correlation analyses comparing relationship between life stage proportions and principal components.** Summary of potential associations for PC1-3 differentially between different life stage proportions inferred with scaden and using *Malaria Cell Atlas* single cell reference sets for *P. knowlesi* (rings, schizonts and trophozoites) and *P. berghei* (gametocytes).

| Upregulated | Protein | Adjusted *p* | LogFC | LogFC SE |
|---|---|---|---|---|
| Severe | unspecified product | 0.033 | 2.879 | 0.612 |
| | high mobility group protein B2, putative | 0.039 | 1.398 | 0.338 |
| | CCR4-NOT transcription complex subunit 1, putative | 0.039 | 1.340 | 0.325 |
| | hypothetical protein, conserved | 0.039 | 1.432 | 0.355 |
| | unspecified product | 0.039 | 2.868 | 0.711 |
| | meiotic recombination protein DMC1, putative | 0.039 | 1.419 | 0.356 |
| | blood-stage antigen 41-3, putative | 0.039 | 1.340 | 0.333 |
| | calpain, putative | 0.039 | 1.814 | 0.453 |
| | AAA family ATPase, putative | 0.039 | 1.095 | 0.276 |
| | erythrocyte membrane-associated antigen, putative | 0.04 | 1.293 | 0.328 |
| | 28 kDa ookinete surface protein, putative | 0.041 | 1.769 | 0.451 |
| | formin 2, putative | 0.044 | 1.139 | 0.296 |
| | subtilisin-like protease 2, putative | 0.044 | 1.113 | 0.289 |
| | LCCL domain-containing protein | 0.046 | 1.377 | 0.366 |
| | reductase, putative | 0.046 | 1.235 | 0.332 |
| Uncomplicated | Protein-L-isoaspartate O-methyltransferase domain-containing protein 1 | 0.033 | -2.021 | 0.435 |
| | Cyclin-dependent kinase 19 | 0.035 | -1.518 | 0.348 |
| | Histone-lysine N-methyltransferase 2C | 0.035 | -1.421 | 0.328 |
| | Protein phosphatase 1B | 0.035 | -1.544 | 0.354 |
| | MORC family CW-type zinc finger protein 3 | 0.038 | -1.973 | 0.464 |
| | SAFB-like transcription modulator | 0.038 | -1.917 | 0.452 |
| | SWI/SNF-related matrix-associated actin-dependent regulator of chromatin subfamily A member 5 | 0.039 | -1.586 | 0.398 |
| | Zinc finger CCCH domain-containing protein 6 | 0.039 | -1.681 | 0.410 |
| | Serine/threonine-protein kinase tousled-like 1 | 0.039 | -1.427 | 0.353 |
| | RNA-binding protein 33 | 0.039 | -1.874 | 0.447 |
| | Zinc finger protein 236 | 0.039 | -1.608 | 0.395 |

| | | | | |
|---|---|---|---|---|
| | Regulator of cell cycle RGCC | 0.039 | -1.878 | 0.466 |
| | Transcription factor Dp-2 | 0.039 | -1.420 | 0.349 |
| | Cyclin-dependent kinase 17 | 0.039 | -1.573 | 0.391 |
| | MAP/microtubule affinity-regulating kinase 3 | 0.039 | -1.594 | 0.402 |

**Supplementary Table 10. Top 15 differentially expressed parasite genes for uncomplicated and severe *P. knowlesi* malaria with brief, tentative biological description.** Genes with significant expression differences (FDR-adjusted $p < 0.05$) are shown, with annotation source indicated. Full results including log2 fold changes, standard errors, all annotations, mutability metrics, and more, for all 130 significant transcripts (112 annotated) are provided in Supplementary File 3.

| Gene | Description | p-adj (voom) | logFC (voom) | Source |
|---|---|---|---|---|
| TRINITY_DN680050_c0_g1 | NA | <0.001 | 0.792 | NA |
| TRINITY_DN112057_c8_g1 | NA | <0.001 | 0.784 | NA |
| TRINITY_DN949463_c0_g1 | NA | <0.001 | 0.723 | NA |
| TRINITY_DN762379_c0_g1 | NA | <0.001 | 0.844 | NA |
| TRINITY_DN47323_c0_g1 | 28 kDa ookinete surface protein, putative | <0.001 | 0.594 | PKA1H1 |
| TRINITY_DN9599_c0_g1 | reductase, putative | <0.001 | 0.536 | PKA1H1 |
| TRINITY_DN1381511_c4_g1 | NA | <0.001 | 0.974 | NA |
| TRINITY_DN299291_c1_g1 | unspecified product | <0.001 | 0.821 | PKA1H1 |
| TRINITY_DN11906_c0_g2 | NA | <0.001 | 0.520 | NA |
| TRINITY_DN14557_c0_g1 | erythrocyte membrane-associated antigen, putative | <0.001 | 0.575 | PKA1H1 |
| TRINITY_DN38550_c1_g1 | NA | <0.001 | 0.740 | NA |
| TRINITY_DN279326_c2_g1 | unspecified product | <0.001 | 0.702 | PKA1H1 |
| TRINITY_DN35871_c0_g1 | meiotic recombination protein DMC1, putative | <0.001 | 0.519 | PKA1H1 |
| TRINITY_DN360328_c22_g1 | NA | <0.001 | 0.620 | NA |
| TRINITY_DN25702_c0_g1 | hypothetical protein, conserved | <0.001 | 0.526 | PKA1H1 |
| TRINITY_DN14177_c0_g1 | AAA family ATPase, putative | <0.001 | 0.534 | PKA1H1 |
| TRINITY_DN99357_c0_g1 | calpain, putative | <0.001 | 0.629 | PKA1H1 |
| TRINITY_DN31815_c0_g1 | conserved Plasmodium protein, unknown function | <0.001 | 0.474 | PKA1H1 |
| TRINITY_DN11069_c0_g1 | erythrocyte membrane-associated antigen, putative | <0.001 | 0.487 | PKA1H1 |

| | | | | |
|---|---|---|---|---|
| TRINITY_DN11835_c0_g1 | conserved Plasmodium protein, unknown function | <0.001 | 0.543 | PKA1H1 |
| TRINITY_DN10352_c0_g1 | conserved Plasmodium protein, unknown function | <0.001 | 0.471 | PKA1H1 |
| TRINITY_DN1116_c0_g1 | conserved Plasmodium protein, unknown function | <0.001 | 0.476 | PKA1H1 |
| TRINITY_DN1350_c0_g1 | conserved Plasmodium protein, unknown function | <0.001 | 0.514 | PKA1H1 |
| TRINITY_DN4936_c0_g1 | blood-stage antigen 41-3, putative | <0.001 | 0.492 | PKA1H1 |
| TRINITY_DN4045_c2_g1 | NA | <0.001 | 0.492 | NA |
| TRINITY_DN16271_c0_g1 | high mobility group protein B2, putative | <0.001 | 0.476 | PKA1H1 |
| TRINITY_DN50536_c0_g1 | subtilisin-like protease 2, putative | <0.001 | 0.506 | PKA1H1 |
| TRINITY_DN10699_c1_g1 | conserved Plasmodium protein, unknown function | <0.001 | 0.547 | PKA1H1 |
| TRINITY_DN18062_c0_g1 | conserved Plasmodium protein, unknown function | <0.001 | 0.547 | PKA1H1 |
| TRINITY_DN19359_c0_g1 | LCCL domain-containing protein | <0.001 | 0.494 | PKA1H1 |

**Supplementary Table 11. Overlapping differentially expressed genes for severity and parasitemia models.** Parasitemia model was performed as described in the methods, except with the continuous $log_{10}$-transformed parasitemia term in place of severity. The limma/voom statistics are presented here, along with the top annotation (prioritisation methods described in methods).

| gene | IDC life stage | Rho |
|---|---|---|
| TRINITY_DN299291_c1_g1 | Gametocyte developing | 0.59 |
| TRINITY_DN949463_c0_g1 | Ring | -0.54 |
| TRINITY_DN949463_c0_g1 | Gametocyte, total | 0.53 |
| TRINITY_DN949463_c0_g1 | Gametocyte, male | 0.53 |
| TRINITY_DN299291_c1_g1 | Gametocyte, male | 0.58 |
| TRINITY_DN949463_c0_g1 | Gametocyte, developing | 0.52 |
| TRINITY_DN299291_c1_g1 | Gametocyte, total | 0.57 |
| TRINITY_DN360328_c22_g1 | Gametocyte, male | 0.56 |
| TRINITY_DN360328_c22_g1 | Gametocyte, total | 0.53 |
| TRINITY_DN299291_c1_g1 | Ring | -0.52 |
| TRINITY_DN360328_c22_g1 | Gametocyte, developing | 0.51 |

**Supplementary Table 12. Life cycle stage associations of differentially expressed transcripts.** Summary of 11 transcripts significantly differentially expressed between severe and uncomplicated *P. knowlesi* infections that are putatively associated with specific life cycle stages predicted with Scaden and *P. knowlesi* (asexual stages) and *P. berghei* (sexual stages) reference data. All associations passed multiple-testing correction (adjusted $p < 0.0001$).

| Wilcoxon rank-sum test | |
|---|---|
| W | 28426 |
| P-value | < 0.001 |

**Supplementary Table 13. Wilcoxon rank-sum test comparing parasitemia between severe and uncomplicated malaria cases.** Parasitemia values were $\log_{10}$-transformed prior to testing.

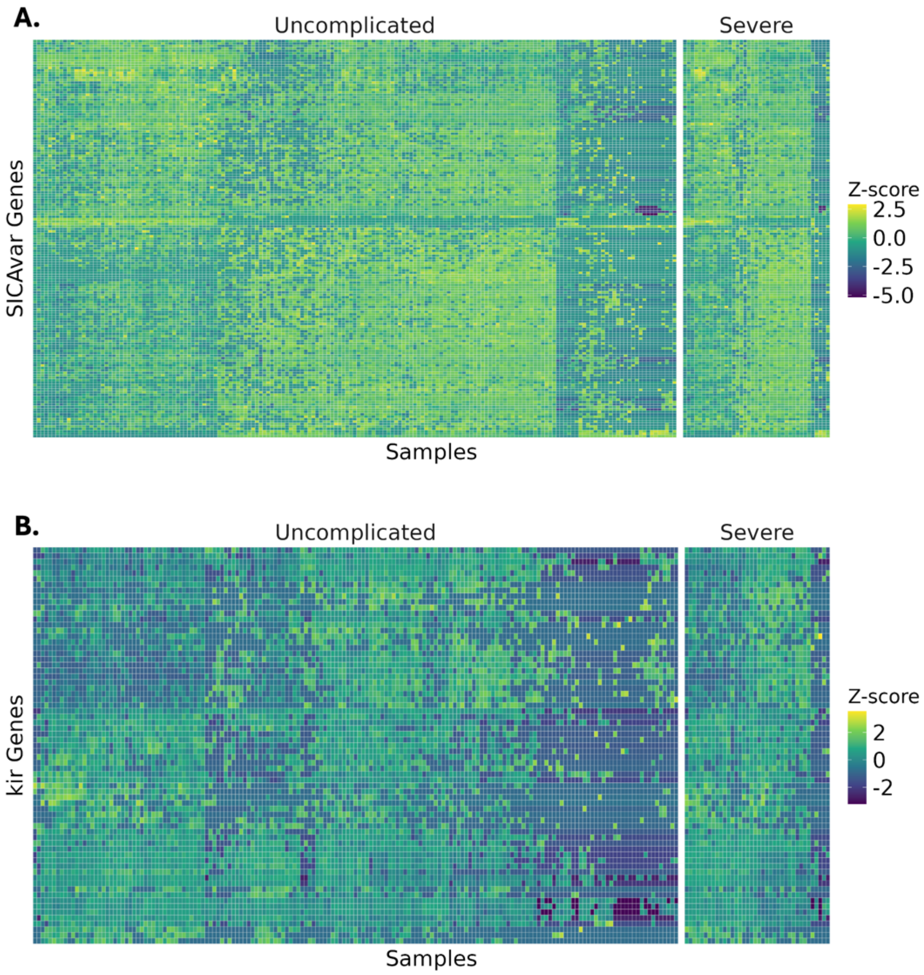


**Supplementary Figure 9. Targeted heatmaps of SICAvar and KIR gene families reveals heterogeneity across families, despite elevated expression in severe malaria.** (A) Sample-level z-score heatmap of filtered SICAvar gene expression across all samples. Columns represent individual patients, and rows represent filtered SICAvar transcripts (logCPM >10 in ≥30% of samples). Z-scores were calculated per transcript across all samples. Both gene-level and grouped-level heatmaps were clustered to highlight expression patterns. (B) Sample-level z-score heatmap of KIR gene expression using the same filtering and normalization approach as panel A. Z-scores were calculated using logCPM-normalized expression values after gene-wise centering and scaling.

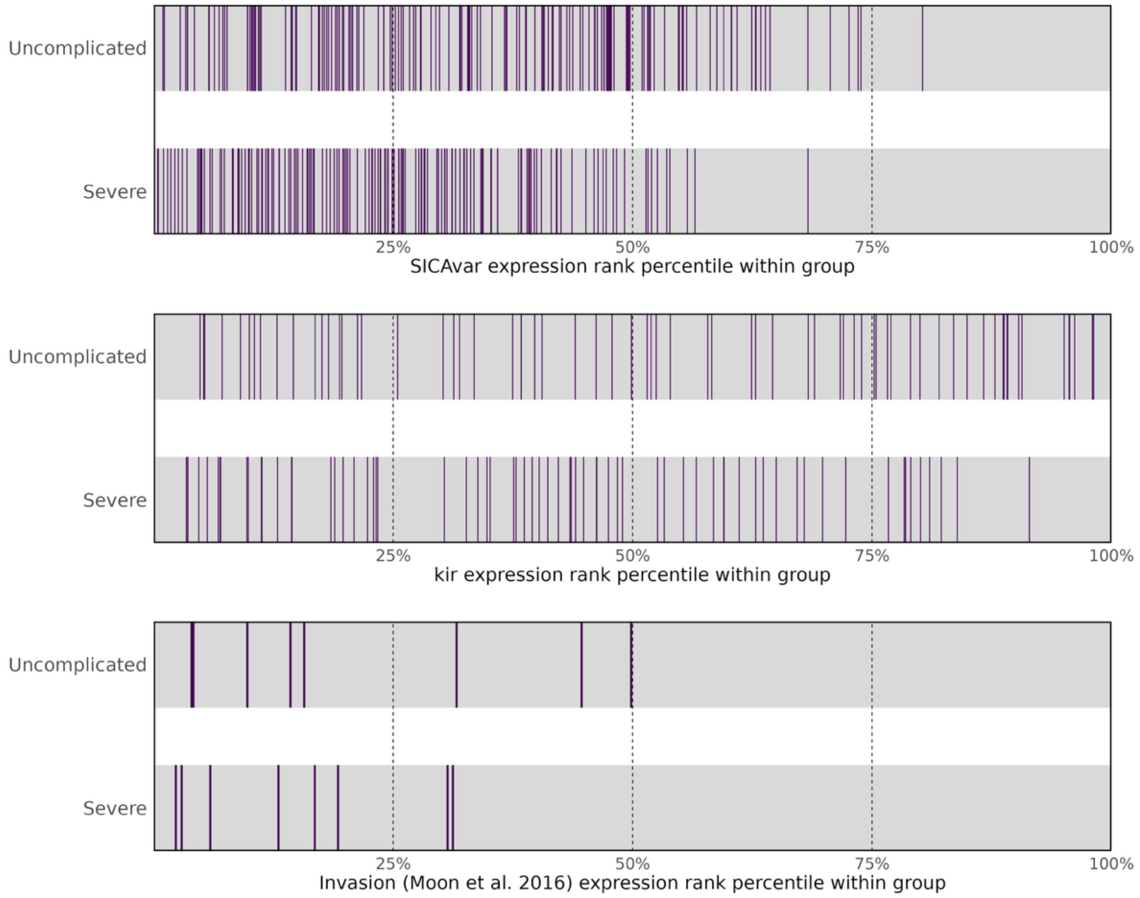


**Supplementary Figure 10. Global expression rank distribution of targeted gene families across clinical severity groups.** Barcode plots show the relative expression rank percentiles of transcripts or genes of interest within the full parasite transcriptome, computed separately for uncomplicated and severe infections. For each group, all expressed transcripts were ranked by mean logCPM expression, and the positions of target features are shown as vertical bars. Top: SICAvar transcripts; middle: kir transcripts; bottom: invasion-related genes curated from Moon et al. (2016), collapsed to gene-level expression prior to ranking. Grey bands represent the full ranked transcriptome, with dashed lines indicating the 25th, 50th, and 75th expression percentiles.

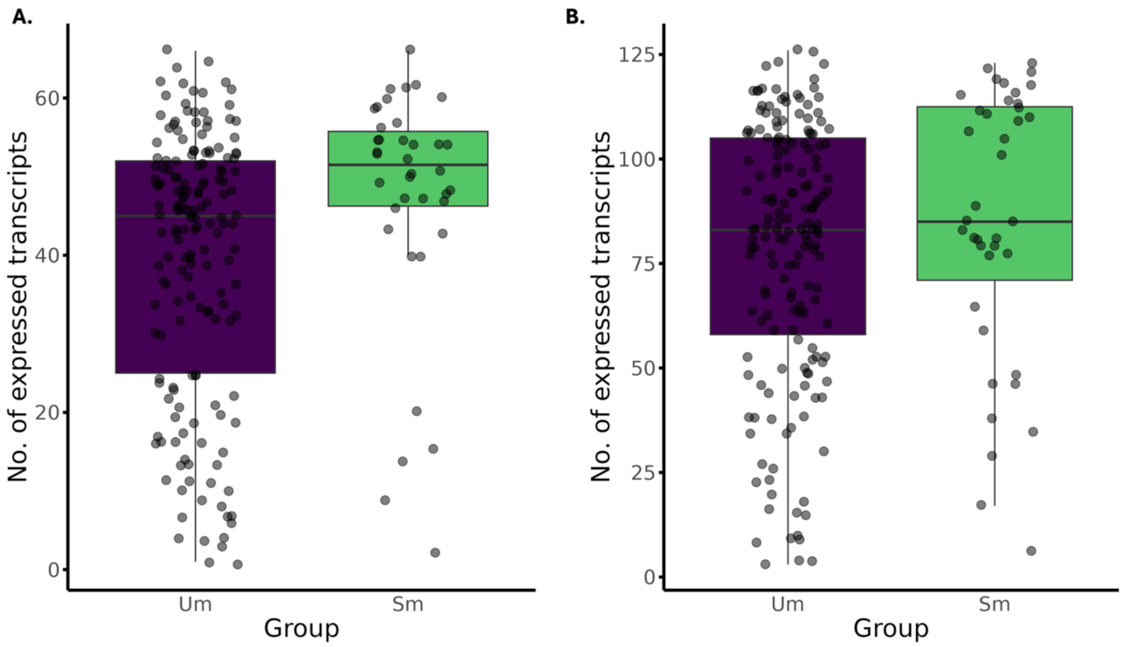


**Supplementary Figure 11. Per-sample kir and SICAvar transcriptional burden in severe and uncomplicated malaria. (A)** Number of kir transcripts expressed per infection in uncomplicated malaria (Um) and severe malaria (Sm) cases. **(B)** Number of SICAvar transcripts expressed per infection in uncomplicated and severe malaria cases. Each point represents an individual infection; boxplots indicate the median and interquartile range, with whiskers extending to 1.5× the IQR. Both gene families show higher transcriptional burden in severe malaria, with kir exhibiting a significantly greater number of expressed transcripts per infection compared with uncomplicated cases (Wilcoxon test; see Results).

| Analysis | Test Type | Result | Adjusted |
|---|---|---|---|
| Fever vs Severe Malaria | Wilcoxon rank-sum test | $p = 0.2311$, $W = 2907.5$ | No |
| Temperature vs Parasitemia | Spearman correlation | $p = 0.16$, $rho = 0.0975405$ | No |

| | | | |
|---|---|---|---|
| Temperature vs Expression of ‘Stress’ Genes (logCPM expression of curated heat-related transcripts [GO terms]) | Spearman correlation per gene | No significant hits. | Yes |

**Supplementary Table 14. Summary of targeted statistical analyses examining the relationship between fever, parasitemia, and heat-responsive gene expression in Plasmodium knowlesi infections.** Gene sets were derived from GO annotations linked to heat stress; both ‘heat’ search in GO terms and curated list from a publication by Zhang et. al. (2). Significance was assessed using non-parametric tests and adjusted for multiple testing where applicable.

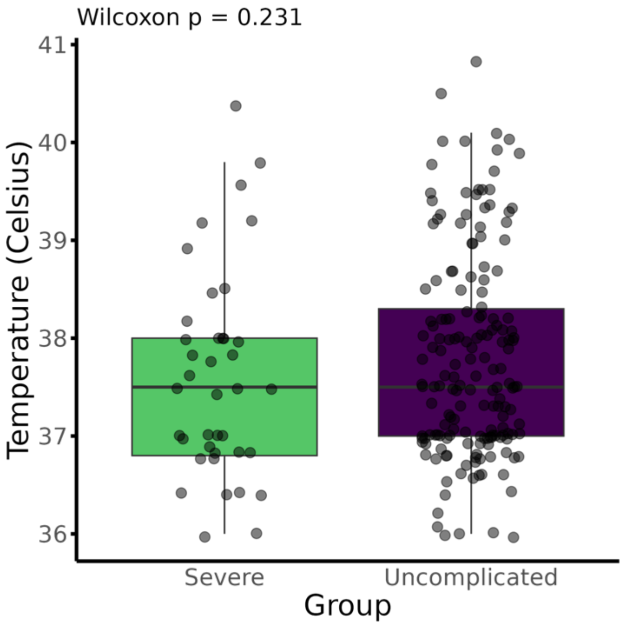


**Supplementary Figure 12. Boxplot comparing the distributions of temperature at admission of patients with severe and uncomplicated malaria.**

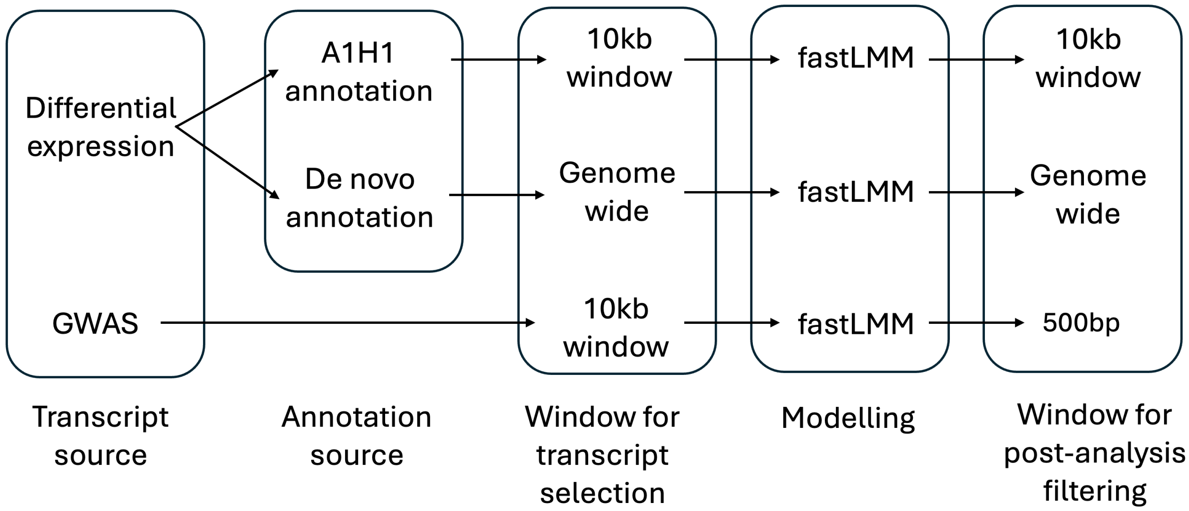


**Supplementary Figure 13. Overview of the integrative framework used to identify expression quantitative trait loci (eQTLs) in Plasmodium knowlesi.** Schematic of the analytical workflow integrating GWAS, differential expression, transcriptome annotation, and multiple statistical modelling approaches to prioritise cis- and trans-eQTL signals. All statistical thresholds and covariates are described in Methods.

1. Howick VM, Russell AJC, Andrews T, Heaton H, Reid AJ, Natarajan K, et al. The Malaria Cell Atlas: Single parasite transcriptomes across the complete Plasmodium life cycle. Science. 2019;365(6455):eaaw2619.

2. Zhang M, Wang C, Oberstaller J, Thomas P, Otto TD, Casandra D, et al. The apicoplast link to fever-survival and artemisinin-resistance in the malaria parasite. Nature Communications. 2021;12(1):4563.
